# Rare bioactive diffusible tau species from Alzheimer brain support both templated misfolding and fibril formation

**DOI:** 10.1101/2025.09.24.678227

**Authors:** Noé Quittot, Dhanush Sivasankaran, Dorothea Böken, Yu Chen, Joshua E. Chun, Anne E. Wiedmer, Victoria Derosla, Mariana B. M. S. Martins, Forest A. Brooks, Georg Meisl, Matthew W. Cotton, Durga G. Arumuganainar, Theresa Connors, Alexandra Melloni, Mathew Frosch, Derek H. Oakley, Lee Makowski, Meni Wanunu, David Klenerman, Bradley T. Hyman

**Affiliations:** Department of Neurology, Massachusetts General Hospital, Boston, MA, USA; Harvard Medical School, Cambridge, MA, USA; UK Dementia Research Institute at the University of Cambridge, Cambridge, UK; Yusuf Hamied Department of Chemistry, University of Cambridge, Cambridge, UK; Department of Bioengineering, Northeastern University, Boston, MA, USA; Department of Physics, Northeastern University, Boston, MA, USA; Technical University of Munich, Munich, Germany; Bioinformatics Program, Northeastern University, Boston, MA, USA; Department of Pathology, Massachusetts General Hospital, Boston, MA, USA; Department of Chemistry and Chemical Biology Northeastern University, Boston, MA, USA

**Keywords:** Alzheimer’s disease, human brain tissue, Tau, oligomer, seed, bioactive species, non-bioactive species, phosphorylations, prion-like mechanism, super resolution microscopy, RT-QuIC

## Abstract

In Alzheimer’s disease, both classical neurofibrillary tangles, and diffusible, aqueous soluble (High Molecular Weight, or HMW) species are able to support templated misfolding. How these tau proteoforms relate is uncertain. Using sequential size exclusion and anion exchange chromatography, we fractionated the HMW tau population and found both seed competent, and seed not competent proteoforms. Super resolution, atomic force, and immunogold electron microscopy confirmed that the size and conformation of both bioactive and non-bioactive tau proteoforms are similar, with dimers, trimers, and tetramers predominating. The presence of surface phosphorylation correlates with seeding capacity. Bioactive tau at fMol concentrations can induce seeding in a reporter cell. The soluble bioactive species support aggregation of a truncated repeat domain tau construct into thioflavin T positive fibrils and retain seeding activity over serial amplification in vitro and in cellulo, whereas non-bioactive oligomeric species do not. Together, these findings indicate that oligomeric assembly is required but not sufficient for seeding; instead, specific biochemical attributes of a rare oligomeric tau subset confer self-propagating, prion-like templated misfolding.

## Introduction

In Alzheimer’s disease (AD), cognitive decline correlates with the hierarchal propagation of tau from medial temporal lobe areas to limbic and association areas^1,2^, consistent with a templated misfolding mechanism of tau seeding and disease progression^3^. Potentially pathological tau is present in a wide variety of tau conformations including classical fibrillar neurofibrillary tangles (NFTs) as well as more recently described diffusible, aqueous soluble, highlighting the structural diversity of the protein^4^. While NFTs were historically considered to be associated with neuronal loss and toxicity, many studies have now shown that intermediate species can also be harmful to the brain^5,6^. In animal models it has been shown that oligomer formation precedes the appearance of NFTs and the spreading is associated with cognitive deficits and neurodegeneration^7–10^. In human AD brains, conformationally specific antibodies detect tau pathology prior to and distinct from classical tangles^11^. Tau oligomer levels are elevated in the absence of NFTs, tau competent seeds are found in synaptically connected networks preceding the formation of NFTs^12,13^, and the amount of tau bioactivity present in the diffusible, aqueous soluble fraction of the brain corresponds with clinical rate of progression in AD^14,15^. While the presence of oligomers is supported by a growing body of evidence, their molecular identity is largely uncertain, and it is unknown how their size and biochemical signature relate to their bioactivity. To date, there is no consensus on how to define tau seeds, particularly in terms of biophysical and biochemical attributes, and whether they are potent amplifiers of structural information.

In an attempt to characterize and identify tau seeds among the numerous varieties of tau species, we reported that high molecular weight (HMW) tau isolated via size exclusion chromatography (SEC) from human brain can be taken up by neurons and induce tau aggregation^16^. This represented <1% of all soluble tau present in the AD cortex. However, this semi-purified fraction, based only on size, is not homogeneous. For example, tau immunoprecipitation methods with monoclonal antibodies directed against various PTMs are not equivalent in removing bioactivity of the HMW samples, suggesting the presence of multiple species^14^. Therefore, in this study we have used a combination of biochemical approaches to fractionate the HMW tau population and revealed the presence of numerous subspecies including both seed competent and non-seed competent species. Moreover, we show that the seed competent species, or bioactive seeds, are retained on an anion exchange column, are sensitive to phosphatase treatment, are oligomeric as imaged with immunoelectron microscopy, super resolution microscopy, and atomic force microscopy, and has phospho-epitopes displayed on the surface. These further purified species can replicate *in vitro,* and daughter seeds generated from the bioactive species induce tau aggregation both *in vitro* and *in cellulo*, whereas daughter seeds derived from non-bioactive oligomeric species remain inactive. These data suggest that seed competence is not dictated by oligomeric state alone and require specific biochemical properties that enable propagation of aggregation over serial passages, consistent with prion-like templated misfolding.

The heterogeneity of aqueous, diffusible tau species—and the absence of consensus biophysical criteria for what constitutes a “seed”—has led to substantial variability in nomenclature across the literature. Here, we use the term “high molecular weight” tau to refer to the characteristic of tau isolated from disease brains as eluting from a size exclusion column in a “high molecular weight” (e.g. several hundred kD) fraction. By contrast, low molecular weight refers to the monomeric tau that elutes from a size exclusion column at the expected molecular weight of tau, and is also referred to as “physiological tau”. We use oligomeric tau to describe small assemblies of tau molecules, not necessarily implying a specific number or specific structure. Fibrils are generally defined as having a β pleated sheet structure, and are observed in Alzheimer tissue as neurofibrillary tangles. Short fibril species are either building blocks or small fragments of the large fibrillar structures that can be observed neuropathologically. The nature of the smallest fragments makes definitive structural determination difficult, in that molecular definition is not possible short of cryoEM, but structures that are not repeated, are not purified, or are not unique are difficult to resolve by cryo-EM. Other biophysical methods, such as those used here including transmission immunoelectron microscopy, atomic force microscopy, and super resolution microscopy do not have sufficient resolution to observe molecular structure directly, but do provide insight into the nature of tau containing assemblies. Because of these limitations, we refer to the material studied here as aqueous diffusible assemblies, and use the term “high molecular weight” to reflect their characteristics on size exclusion columns. While fibrils were not observed in any of our experimental preparations, the possibility that there are some short fibrils cannot be definitively ruled out.

## Methods

### Human tissue and data collection

Nine human participants with AD were selected from the Massachusetts Alzheimer’s Disease Research Center Longitudinal Cohort study on the basis of the following criteria: (1) clinically diagnosed with dementia due to probable AD; (2) cognitive status assessed at least three times by a neurologist at the Massachusetts General Hospital (MGH) Memory Clinic Unit and scored using CDR-SOB; (3) diagnosis of AD confirmed postmortem by an MGH neuropathologist; (4) Braak Neurofibrillary tangles (NFTs) stage V or VI as determined by the location of NFTs with a total tau immunostaining^17^; and (5) concurrent pathologies considered to be clinically non-significant. Age of onset, age at death, postmortem interval and sex were also collected and are listed in Supplementary Table 1. Autopsy tissue from human brains were collected at MGH, with informed consent and approval of local institutional review boards. Human brains were processed as previously described^13^. Briefly, all brains were separated into 2 hemispheres, one of which was initially sliced coronally at the time of autopsy and 1-cm-thick slabs were immediately flash frozen and stored at −80 °C. Approximately 5 g of frontal cortex was dissected out of the frozen brain section corresponding to Brodmann Area 8 and kept at −80 °C until homogenization.

### Tissue brain homogenization

Frozen human tissue (5g) was thawed on wet ice, and gray matter was dissected and then immediately homogenized in PBS containing 1X protease and phosphatase inhibitors (Cell Signaling, #5872) + 2 µM Trichostatin A (Sigma, #T8552) in a 15-ml glass Dounce homogenizer. The tissue was Dounce homogenized with 30 up and down strokes on ice by hand. The homogenate was centrifuged at 10,000g for 10 min at 4 °C. The supernatant was collected and aliquoted to avoid excessive freeze–thaw cycles.

### Tau seed enrichment

Each chromatographic step was performed on a fast protein liquid chromatography (FPLC) apparatus (ÄKTA Pure 25L, GE Healthcare) at 4°C. All buffers were filtered through a 0.2-μm membrane filter and cooled down to 4°C before use.

### Size exclusion chromatography (SEC)

5 mL of brain PBS extracts were separated by SEC on a single HiLoad 16/600 Superdex 200 pg column (no. 28989335, Cytiva) in Tris 20 mM, pH 8.6 at a flow rate of 1 ml.min^−1^. Fractions of 2 mL were retrieved and analyzed by western blots, and subjected to the *in vitro* seeding assay. Fractions containing bioactive HMW tau species were pooled.

### Anion exchange chromatography (AIEX)

5mL of HMW tau species were enriched via a high-resolution and strong anion exchange chromatography medium, consisting of non-porous 3 µm monodisperse beads, using a Mini Q 4.6/50 PE column (no. 17517701, Cytiva). HMW tau species were captured on the resin with a starting buffer of Tris 20 mM, pH 8.6 and eluted by applying a 20 mL linear gradient, or step-wise elution, with Tris 20 mM + 4 M NaCl, pH 8.6 at a flow rate of 0.25 ml.min^−1^. Fractions of 0.5 mL were retrieved and analyzed by dot blot and the *in vitro* seeding assay. Fractions containing non-bioactive and bioactive HMW tau species were pooled separately.

### Desalting column

Following AIEX, non-bioactive and bioactive HMW tau species were buffer exchanged on a HiTrap desalting column (no. 29048684, Cytiva) to phosphate-buffer saline (PBS) at flow rate of 5 ml.min^−1^. A minimum threshold of 5 mAU was set to start/stop collection. Conductivity was monitored to evaluate the buffer exchange from a low salt to a high salt buffer.

### Western Blots and Silver Stain

NuPAGE LDS Sample Buffer (ThermoFisher, NP0007) and NuPAGE Sample Reducing Agent (ThermoFisher, NP0009) were added to tau seeds, boiled for 5 min at 95 °C and run on 4–12% Bis-Tris SDS–PAGE (Invitrogen) in MOPS buffer (Invitrogen). The gels were stained with silver stain following manufacturer instructions (Pierce^TM^ Silver Stain Kit, no. 24612) or transferred to a nitrocellulose membrane (NP0324BOX) using an Invitrogen™ iBlot™ 2 Gel Transfer Device (IB21001). Blots were incubated with blocking buffer (Intercept TBS blocking buffer, Li-Cor, no. 927-60001) for 30 minutes followed by incubation with a monoclonal rabbit anti-pan-tau 1:1,000 (D5D8N, no. 43894, from Cell Signaling) overnight at 4 °C or 1h at room temperature. Blots were washed three times for 10 min with TBS-T and incubated with secondary anti-rabbit IRDye680 or IRDye800 (1:10,000) for 1 h at room temperature and imaged on the Odyssey infrared imaging system (Li-Cor).

### Dot Blots

30 to 100 µL of fractions were loaded on a nitrocellulose membrane using a 96-well dot blot apparatus (Bio-Rad, no. 1706545) and incubated for 30 min at room temperature before proceeding to vacuum filtration. Blots were then processed as described in the western blot section. Dot blot intensities were quantified by by densitometric analysis in Image J (Fiji).

### Two-dimensional gel electrophoresis (2D gel)

30 µL of the sample was mixed with 1.4 µL Blue Bromophenol 1% and 108.6 µL rehydration buffer (BioRad #1632106). The mix was loaded in a 7 cm running chamber of a six-enclosed channels cassette (ThermoFisher, ZM0003). A 7-cm IPG strip with a non-linear gradient pH ranging from 3 to 10 (ThermoFisher, ZM0018) was added to the running chamber containing the sample mix. The cassette was sealed and incubated for 1 hour at room temperature. For the first dimension, the cassette was placed in the IPG runner mini cell (ThermoFisher, ZM0001), and the outer chamber was filled with 600 mL of deionized water. The isoelectric focusing was performed using ZOOM™ Dual Power Supply (100-120 VAC 47 - 60 Hz) (ZP10001) with the following sequence: 1) 175 V for 30 min; 2) 175 to 2000 V over 90 min; 3) 2000 V for 60 min. For the second dimension, strips were incubated with NuPAGE LDS Sample Buffer (ThermoFisher, NP0007) and NuPAGE Sample Reducing Agent (ThermoFisher, NP0009) using an equilibration tray (ThermoFisher, ZM0007) at room temperature for 15 min. Strips were then loaded on NuPAGE™ Bis-Tris Mini Protein Gels, 4–12%, 1.0–1.5 mm (NP0330BOX) and sealed by adding 0.5% agarose mixed with MOPS running buffer. Electrophoresis is performed in two steps: 1) 100 V for 15 min followed by 2) 175 V for 60 min. Gels were then uncast and transferred to a nitrocellulose membrane as previously described.

### Dephosphorylation

HMW tau was subjected to dephosphorylation using Lambda Protein Phosphatase (LPP) kit (New England Biolabs (NEB), #P0753S). Two reaction mixes were prepared: one containing LPP at concentration of 800 units/mL and another without the enzyme. Each reaction was performed with NEB buffer for Protein MetalloPhosphatases, 1 mM MnCl₂, 1X Protease Inhibitor Cocktail. Both reaction mixes were incubated at 30 °C for 2 hours. To inactivate phosphatase, samples were heated at 65 °C for 1 hour. Samples were then centrifuged at 15 000g for 10 minutes at 4 °C to pellet precipitated enzymes. The supernatants were collected and stored at –20 °C for further analysis.

### Atomic Force Microscopy

To characterize the morphology of the bioactive and non-bioactive tau species, 5µL protein samples were mixed with 5µL phosphate-buffered saline (PBS). This mixture was subsequently deposited onto a freshly cleaved mica substrate and allowed to incubate for 10 minutes. Following incubation, the mica surface was rinsed with deionized water (DI) to remove unbound proteins and then dried under a gentle stream of nitrogen. The dried protein samples were imaged were imaged utilizing a MultiMode 8 AFM and Nanoscope V controller (Bruker, Billerica, MA, USA) performed in ScanAsyst in air mode using a ScanAsyst-air silicon tip (Bruker). The collected imaging data were processed using NanoScope Analysis software (Bruker) and further analyzed by Gwyddion software.

### Single-molecule pull-down (SiMPull)

Single-molecule pull-down (SiMPull) super-resolution microscopy was performed as previously described^18^ with slight modifications. Briefly, passivated PEGylated glass coverslips were incubated with biotinylated capture antibody (10 nM in blocking solution: 1mg/mL BSA in PBS) for 10 minutes. This was followed by washing twice with PBST (0.05% Tween-20 in PBS) and once with 1% Tween-20 in PBS and then 10 minutes incubation in blocking solution. After a PBST wash, 10µL of biological samples (1:2 dilution in PBS) was added and left to incubate for 1 hour at room temperature, followed by washing in PBST twice and 1% Tween-20 and 30 minutes incubation in blocking solution. Detection antibody covalently labelled with a docking strand (DBCO TEG-AAACCACCACCACCACCACCACCACCACCACCACCA) for DNA PAINT was incubated (3nM in 0.1 mg/mL BSA in PBS) for 15 minutes, followed by washing in PBST twice, 1% Tween-20 and PBS. Last, 3µL imaging strands (1 nM in PBS, TGGTGGT-atto655) were added before the gasket was sealed with a clean coverslip.

### Super resolution image acquisition and analysis

Imaging was performed on a home-built total internal reflection fluorescence (TIRF) microscope, comprising an inverted Ti-2 Eclipse microscope body (Nikon) with a 1.49 N.A., 60x TIRF objective (Apo TIRF, Nikon) and a perfect focus system. Images were acquired for 8000 frames of 100 ms exposure using a 638 nm laser (Cobolt 06-MLD-638, HÜBNER) and were collected in a grid using an automated script (Micro-Manager) to avoid any bias in the selection of the field of views. Super-resolution images were reconstructed and analyzed as described in Böken *et al.*, 2024^18^. Briefly, localizations were identified and fit using ComDet then corrected for microscope drift using the inbuilt implementation of redundant cross-correlation. Localizations were filtered for precision <30 nm before DBSCAN clustering was performed using the scitkit-learn package with permissive parameters (radius of 0.3 and minimum density of 5). The skeletonized length of each aggregate was measured whereby the length of each aggregate is reported as the summed branch distance. Finally, super-resolved images were rendered using the inbuilt Picasso functionality.

### Single Molecule Array (SIMOA)

Simoa assays were developed using the Quanterix HomeBrew set up. Bead conjugation was performed as per the manufacturer’s instructions using Quanterix reagents. Briefly, 488 dye-coated paramagnetic carboxylated beads were activated with EDC at 4°C, washed, and subsequently conjugated to antibody, followed by blocking of the beads. The antibody coating efficiency was determined by measuring the residual antibody concentration in the solution and supernatant of the first wash. Beads were pelleted and stored in Bead Diluent Buffer at 4 °C. Simoa plates were prepared in a “3-step-assay” following the Quanterix protocol. Samples were diluted in tau 2.0 sample diluent, and added to a Simoa 96-well plate (100 µL). Approximately 500k antibody-conjugated beads were added to each well and incubated (30°C, 800 rpm, 30 min). All washing steps were performed using the automated 3-step-assay program on the Simoa Microplate Washer. After three automated washes with Simoa wash buffer A, each well was incubated with 100 µL biotinylated detection antibody (0.3 µg/mL in Detector/Sample diluent, 30°C, 800 rpm, 10 min), followed by three further washes with Simoa wash buffer A. SBG (streptavidin β-galactosidase) enzyme was added to each well (150 pM in SBG Dilution buffer and incubated (30°C, 800 rpm, 10 min), followed by three washes with Simoa wash buffer A and two washes with Simoa Wash Buffer B. The beads were left to dry for 10 min and processed on the Quanterix SR-X™ Instrument using RGP as substrate. The readout is determined as the fraction of beads with enzyme activity (*f*_ON_), which calculates the average enzymes per bead (AEB) assuming Poisson statistics. For aggregate-specific assays, any sample or calibrant concentrations with *f*_ON_ > 0.7 was avoided. Lysate from cells overexpressing tau aggregates was used as a calibration standard^19^. After obtaining the AEB values at each calibration level, a four-parameter logistic (4PL) curve was fitted.

### Negative-stain electron microscopy

For negative stain electron microscopy, 3 µL of the desired sample was applied to a 200 mesh Cu Formvar/Carbon grid and incubated for two minutes before the excess liquid was wicked dry and the grid was washed with 3 µL of water. The grid was then wicked dry again and 3 µL of 2% uranyl acetate was applied for 90 seconds before being wicked and allowed to dry. Images were recorded using a JEOL 1011 transmission electron microscope.

### Immunogold labeling for electron microscopy and image analysis

Sample application followed the same steps as outlined for negative-stain electron microscopy. Then, the grid was placed sample side-down onto a drop (15-20 µL) of blocking buffer (PBS + 0.1% gelatin) on a piece of parafilm. Parafilm is placed on a platform in a water dish to maintain humidity. The grid was then left in blocking buffer for 10 minutes. Then, the grid was placed onto a drop of diluted primary antibody (in blocking buffer) for 3 hours. The grid was then washed by being transferred to a drop of blocking buffer for 3 minutes. This was repeated for a total of 4 washes. The grid was then transferred to a drop of undiluted gold conjugated secondary antibody for 1 hour. The grid was subsequently washed by being transferred to a drop of PBS for 3 minutes. This was repeated for a total of 2 PBS washes, and 2 water washes. If double-labeling, grid was placed into a new primary antibody and underwent the following steps (a different size gold particle-conjugated secondary antibody was used). Once the final water wash was wicked away, 3 µL of 2% uranyl acetate was applied for 90 seconds, then wicked dry. Images were recorded using a JEOL 1011 transmission electron microscope. Histograms of distances between gold particles were generated by determining the (x,y) coordinates of the center of each gold particle in an electron micrograph using the ‘Analyze Particles’ utility in ImageJ. Additional processing utilized a simple Python script that (i) calculated the distance between each gold particle pair; (ii) Calculated a raw histogram by binning the inter-particle distances; (iii) divided the number of particles in each bin by the mean gold-gold distance of the corresponding bin. This produced a plot of number density (relative frequency) of particles at each distance. This number density was then normalized to facilitate comparison of results from different samples. The resulting ‘histograms’ thereby provide a measure of how the presence of a gold particle influences the number density of gold particles in its vicinity.

### RT-QuIC assay

Real-time Quaking-Induced Conversion (RT-QuIC) assays were performed by incubating biological samples with two different tau substrates. 1) A Tau fragment corresponding to the AD core, residues 306–378, was used at a final concentration of 10 µM in an assembly buffer composed of 100 mM NaCl, 10 mM HEPES, and 1 mM TCEP, as described in Carlomagno *et al.*, 2021^20^. 2) Recombinant tau isoforms 0N4R and 0N3R were combined at a 2:3 ratio to a total protein concentration of 10 µM in PBS. Biological samples were added at 2 µL per well and incubated with each substrate mixture. Thioflavin T (ThT) was included in each reaction to a final concentration of 25 µM. Reactions were carried out in 40 µL total volume, in triplicate, in sterile, sealed 384-well black-walled, clear-bottom, nonbinding microplates (Corning). Kinetic assays were conducted at 37 °C using a FLUOstar Omega plate reader with continuous orbital shaking at 200 rpm. ThT fluorescence (excitation: 440 nm; emission: 485 nm) was measured every 15 minutes from the bottom of the plate to monitor fibrillization. In separate wells, conditions were replicated without ThT used for negative stain electron microscopy and second run of seeds amplification.

Kinetic data from triplicates were averaged and the lag time (t_lag_), defined as the time before any detectable amyloid forms, was determined with a Boltzmann sigmoidal curve,

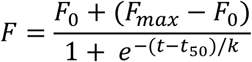

where t_50_ is the time to half-maximum intensity, k is the apparent first-order rate constant, and F_0_ and F_max_ are the minimum and maximum fluorescence intensities, respectively. The lag time was then calculated using the formula t_lag_ = t_50_ − 2k.

### Seeding assay

#### In vitro tau seeding FRET-biosensor assay

The *in vitro* seeding assay has been previously described and widely characterized^21^. Briefly, the Tau RD P301S FRET Biosensor (ATCC CRL-3275) cells stably expressing the repeat domain of tau with the p.P301S mutation conjugated to either cyan fluorescent protein (CFP) or yellow fluorescent protein (YFP) (TauRD-P301S-CFP/YFP) were cultured at 37 °C, 5% CO2 in DMEM, 10% vol/vol FBS, 0.5% vol/vol penicillin–streptomycin. Cells were plated on Costar Black, clear-bottom 96-well plates (previously coated with 1:20 poly-D-lysine) at a density of 20,000 cells per well. The same volume of brain extracts (2.67 μl per well) was then incubated with Lipofectamine 2000 (Invitrogen, final concentration 1% vol/vol) in opti-MEM (final volume of 50 μl per well) for 10 min at room temperature before being added to the cells. Each condition was applied in triplicate. Tau seeding was subsequently analyzed using flow cytometry.

#### Flow-cytometry seeding analysis

After 24 h, medium was removed, and 50 μl trypsin 1× was added for 7 min at 37 °C. Chilled DMEM + 10% FBS (150 μl) was added to the trypsin, and cells were transferred to 96-well U-bottom plates (Corning). Cells were pelleted at 500g, resuspended in freshly made 2% vol/vol paraformaldehyde in PBS (Electron Microscopy Services) for 10 min at room temperature in the dark, and pelleted at 500g. Cells were resuspended in chilled PBS and run on the MACSQuant VYB (Miltenyi) flow cytometer. CFP and FRET were both measured by exciting the cells using the 405-nm laser and reading fluorescence emission at the 405/50-nm and 525/50-nm filters, respectively. To quantify the FRET signal, a bivariate plot of FRET versus the CFP donor was generated, and cells that received control brain extract alone were used to identify the FRET-negative population. Using this gate, the tau seeding value for each well was calculated by multiplying the percentage of FRET-positive cells by the median fluorescence intensity of that FRET-positive population. We analyzed 40,000 events per well. Data were analyzed using the MACSQuantify software (Miltenyi).

#### Fitting of single hit dynamics

Fitting a one hit model to the seeding titration data shows good agreement. The one hit model applies to systems where a single seeding event is required to trigger the production of daughter seeds. A system with a low seeding efficiency can still show one hit dynamics, however a much larger concentration of seed will be needed to see the same response if the seeding efficiency is lower. Whether a system shows one hit dynamics or not will depend on the seeds and the cells/system that is being treated. We fit the following functional form to the data that calculates the number of positive cells given a specific concentration of seeds,

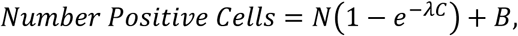

where *N* is the total number of cells, λ is the seeding efficiency and *C* is the seed concentration. The final term, *B*, is an additional baseline of cells that would test positive independent of seed. We set *B* = 3 based on the positive baseline observed in control experiments. For the bioactive species, the best fit for the data set is given when *N* = 950 and the seeding efficiency λ = 1.3 × 10^−4^ fM^-1^. We use the same baseline and total number of cells to estimate the seeding efficiency in the non-bioactive data. There is no pronounced dose response in the non-bioactive data; most seed concentrations give essentially the same number of positive cells. We therefore only consider the highest three seeding concentrations where a dose-response may be emerging. This gives the non-bioactive seeding efficiency as λ = 5.3 × 10^−6^ fM^-1^. As even at those highest three points, the response to seeding is weak compared to background, this should be interpreted as an upper bound on the seeding efficiency of the non-bioactive species.

#### Antibodies

D5D8N and HT7 are both monoclonal antibodies directed against a central tau epitope (∼residue 160) and detect all six canonical CNS tau isoforms and hence are referred as pan-tau antibody; however, it may not detect truncated fragments lacking this region.

AT8 is a monoclonal antibody directed against phosphoresidues Ser 202 and Thr 205.

pT181, pT217, pT231, pS262, pS356, pS396, pS404 are monoclonal antibodies directed against indicated residues (S: Serine, T: Threonine) at the indicated position.

PHF1 is a monoclonal antibody directed against phosphoresidues pS396 and pS404.

## Results

### High-molecular-weight tau contains various subspecies, with the phosphorylated forms being the most seeding-competent

Initial analysis by two-dimensional gel electrophoresis revealed that HMW tau isolated from nine human AD brains consisted of a heterogeneous population of molecules with isoelectric point (pI) values ranging from 5 to 8.5, indicating a mix of modified and unmodified tau species (Supplementary Figure 1). Given that phosphorylation is the predominant modification, we hypothesized that its removal would significantly impact the net charge and seeding activity of HMW tau. To test this, we treated HMW tau with lambda protein phosphatase (Lambda PP) to assess whether we could reduce the seeding activity (Figure 1A). Dephosphorylation significantly reduced tau’s bioactivity, as measured by the FRET reporter HEK cell assay^21^ (Figure 1B). This finding aligns with prior studies showing that dephosphorylation diminishes tau’s seeding competency^16,22^, and with our previous observation of a positive correlation between the extent of post-translational modifications (PTMs) and the bioactivity of HMW tau^23^. Dot blot analysis confirmed the loss of immunoreactivity for the tau phospho-specific AT8 antibody, while tau levels assessed with pan-Tau antibody remained unchanged (Figure 1D, E). In summary, these results indicate that the HMW tau population includes a mix of modified molecules, with highly charged, phosphorylated species being the most bioactive.

**Figure 1.**
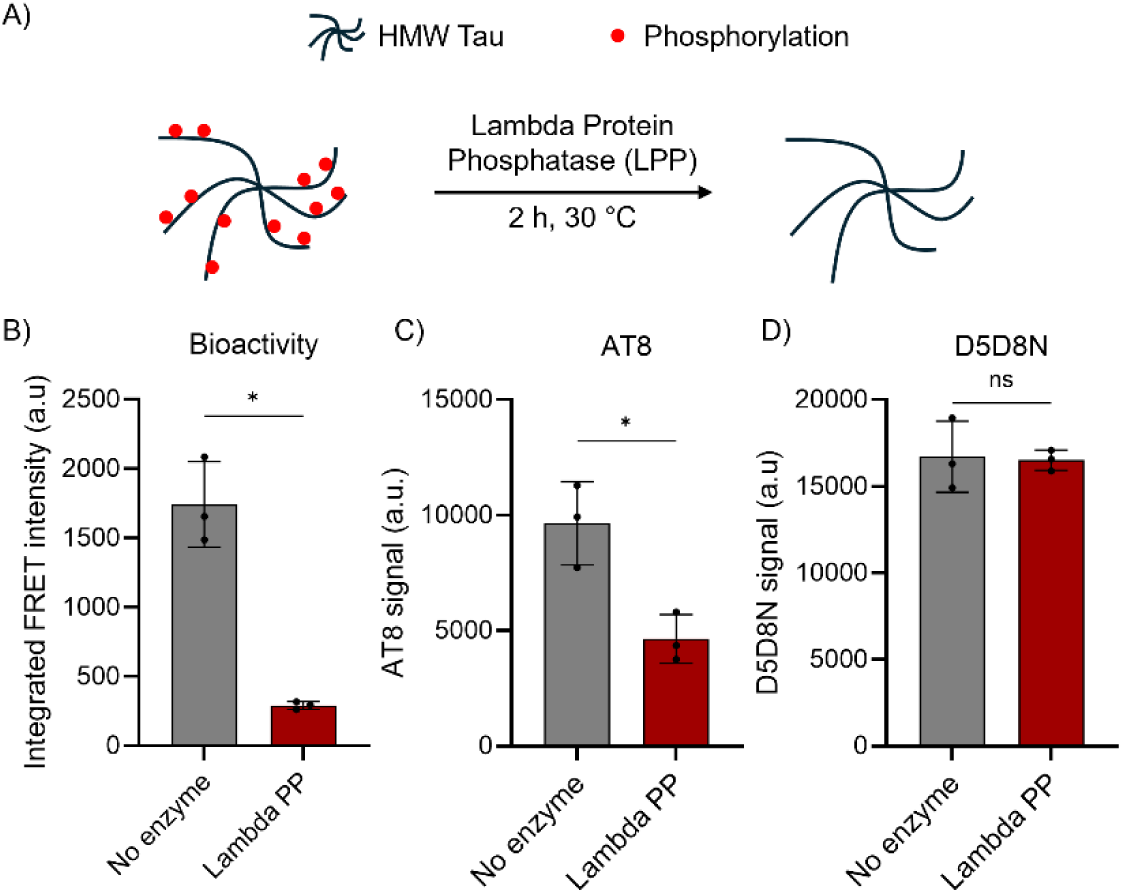
Dephosphorylation of human-derived HMW tau species decreases bioactivity. Schematic representation of HMW tau dephosphorylation by the Lambda Protein Phosphatase (LPP). Seeding activity assay using FRET reporter HEK cell; the integrated FRET intensity was quantified by flow cytometry. Data represents mean ± S.D. of three individual experiments performed in triplicate with HMW Tau isolated from the AD case 1971. Results were analyzed using the paired student t-test, * for p < 0.05 and ns for non-significant. C), D) Dot blot quantification of HMW tau, originating from the same case as panel B, treated or not with Lambda PP and probed with the anti-phospho tau AT8 or the pan-tau antibody D5D8N.

### Both non-bioactive and bioactive species are found in the HMW tau population

Given the variation in apparent pI of tau species from the early SEC fractions observed by 2D gel, we further separated the HMW fraction using a technique sensitive to variations in surface charge. Considering that phosphorylation adds two negative charges at physiological pH, while acetylation or ubiquitination neutralizes lysine side chains, we hypothesized that more negatively charged modified and most bioactive tau species will interact more strongly with the resin and be eluted in later fractions. This strategy has proven to be efficient in separating unmodified and phosphorylated tau produced in yeast and SF9 insect cells ^24,25^. Therefore, following SEC, the collected HMW Tau was applied to an anion exchange column. Elution profiles for Tau and phosphorylated Tau were then determined under native conditions using dot blot analysis, with the pan-Tau antibody D5D8N and the phospho-specific antibody AT8, respectively. The corresponding seeding activity of each eluted fraction was determined with the previously mentioned cell-based assay. Interestingly, we observed that tau was eluted in nearly all the fractions along the linear elution gradient, confirming the presence of multiple tau species varying in their binding capacity to the resin, suggesting species with varying overall net charge and/or conformations (Figure 2A). Further analysis revealed that 72% of the HMW tau eluted consisted of AT8-positive species, while only 44% of the HMW tau eluted were bioactive species. Intriguingly, while every fraction containing the bioactive species was AT8 positive, we also observed non-bioactive AT8 positive species, accounting for 28% of the HMW tau eluted. As expected, the bioactive AT8 positive species were eluted later compared to the AT8 negative species. Importantly, we identified the same pattern of elution of non-bioactive and bioactive species for 9 AD cases with the exact same cut-off for the separation of the two species, with bioactive species eluting at a NaCl concentration of 450 mM (Figure 2A, Supplemental Figure S3). This consistency among cases was surprising considering the case-to-case variations in PTMs as noted in our prior mass spectroscopy analysis using tissue from these same cases^23^.

**Figure 2.**
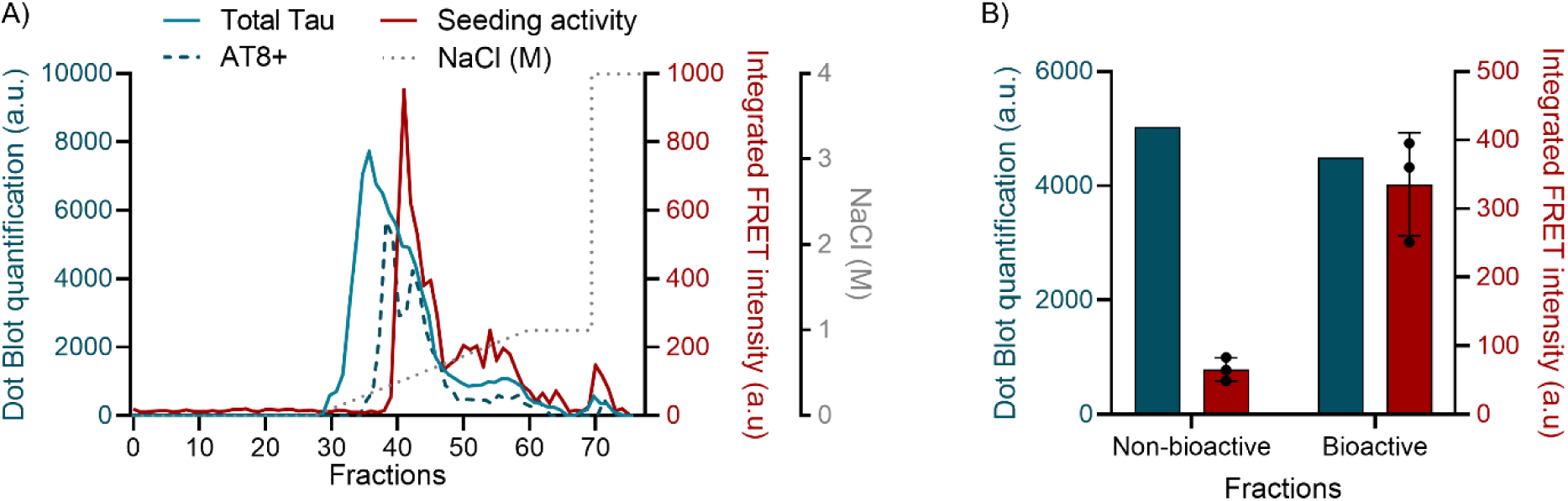
HMW tau fractionation by charge reveals seeding competent and inert species. HMW tau fractionation originating from the AD case 1971 via anion exchange chromatography (AIEX) with increasing NaCl concentration (gray dashed line). Tau and phosphorylated tau are measured in each fraction by dot blot quantification using the pan-Tau antibody D5D8N (blue line) and AT8 antibody (dashed blue line). Seeding activity of each fraction is determined using the FRET reporter HEK cell assay in triplicate and measured by flow cytometry (red line). Data from one seeding assay, out of three, is plotted as representative of the bioactivity profile. B) Separation of non-bioactive and bioactive HMW tau species, originating from the AD case 1971 as in panel A, with AIEX via a step wise elution of NaCl. First-step elution is performed with 0.3M NaCl followed by a second step of 4M NaCl. Non-bioactive and Bioactive fractions are desalted into PBS before tau quantification on dot blot by the pan-Tau antibody D5D8N (blue) and seeding activity assay measured by flow cytometry (red). Seeding assay data represent mean ± S.D. of three individual experiments performed in triplicate, using non-bioactive and bioactive species isolated from AD case 1971.

The linear gradient of elution separates the HMW tau population into multiple fractions, but mainly two species are isolated, the non-bioactive and bioactive species. Therefore, considering the clear cut-off between the non-bioactive and bioactive species, we performed a stepwise elution to separate both species and collected each species within one fraction of 0.5 mL of buffer. In this strategy, both species are retained on the AIEX column in Tris 20 mM pH8.6 as previously described, but for the elution, non-bioactive species are eluted with a constant salt concentration of 0.3 M of NaCl and bioactive species are then eluted with 4M of NaCl. Both fractions were then desalted into PBS. First, we evaluated the tau quantity in the fractions corresponding to the two steps of elution by dot blot in native condition and we observed a similar tau amount eluted in each step (Figure 2B). We then validated the separation of non-bioactive vs bioactive tau species with a seeding assay showing a low bioactivity in the first step of elution comparatively to the high seeding activity in the second step of elution, independently of the tau concentration (Figure 2B). We next asked if the anion exchange chromatography separates two forms of tau or “peels” loosely bound tau from an aggregated form of tau. To test this hypothesis, we injected both species into a SEC column to evaluate the size and the associated seeding activity. Strikingly, the bioactive species contain only HMW tau species, while the non-bioactive species contain a mixture of low and high molecular weight tau (Supplemental Figure S4). Overall, these results suggest that the HMW tau population contains multiple subspecies including non-bioactive and bioactive species that both co-elute from SEC in the HMW fractions but differ by their retention time on an anion exchange column. We hypothesize that this behavior is due to a difference in the number of charges, or a variety of conformations, or both.

### Both bioactive and non-bioactive HMW tau species consist of small diffusible aggregates

We showed that the non-bioactive and bioactive species maintain their aggregate conformation after AIEX and are consistently found in every AD case tested, hence we used one representative AD case (1971) to carry the following biophysical and biochemical experiments. Given the ongoing debate in the field regarding whether tau bioactive species are insoluble fibrils or small diffusible aggregates^26,27^, we carried out atomic force microscopy (AFM) of the samples deposited on freshly-cleaved mica surfaces (representative AFM micrographs are shown in Figure 3A). The images clearly show the absence of fibrillar structures, with mostly globular shapes, although these qualitatively differ for the bioactive and non-bioactive samples. Particle size analysis of the AFM images of the bioactive and non-bioactive fractions allowed us to extract two critical parameters, namely the equivalent radius (r_eq_) and the maximum particle height (Z_max_), the distributions of which are shown in Figure 3B. Interestingly, we find a mono-exponential distribution of r_eq_ values for both samples, with a decay constant of 2.65 nm and 3.72 nm for bioactive and non-bioactive, respectively. However, analysis of Z_max_ values reveals two populations for each of the samples, with a consistent cutoff maximum height of ∼5 nm for both populations. Fitting the distributions to bi-exponential functions, we find that the fraction of higher Z particles, *F*_2_, is greater for the bioactive particles (23*±1.16*%) than for the non-bioactive particles (2*±0.01%*), as calculated using *F*_2_ = *a*_2_*z*_2_/(*a*_1_*z*_1_+*a*_2_*z*_2_), where *a* are the amplitudes and *z* are the decay constants obtained from the bi-exponential fits. In summary, AFM did not show any fibrillar structures for either bioactive or non-bioactive particles, and the observed particle size ranges are smaller than the sizes typically reported for fibrillar tau aggregates^28,29^. To note, using human brain lysate from a control case, we observed similar size particles with r_eq_ values ranging from 3 to 40 nm (Supplemental Figure S4).

**Figure 3.**
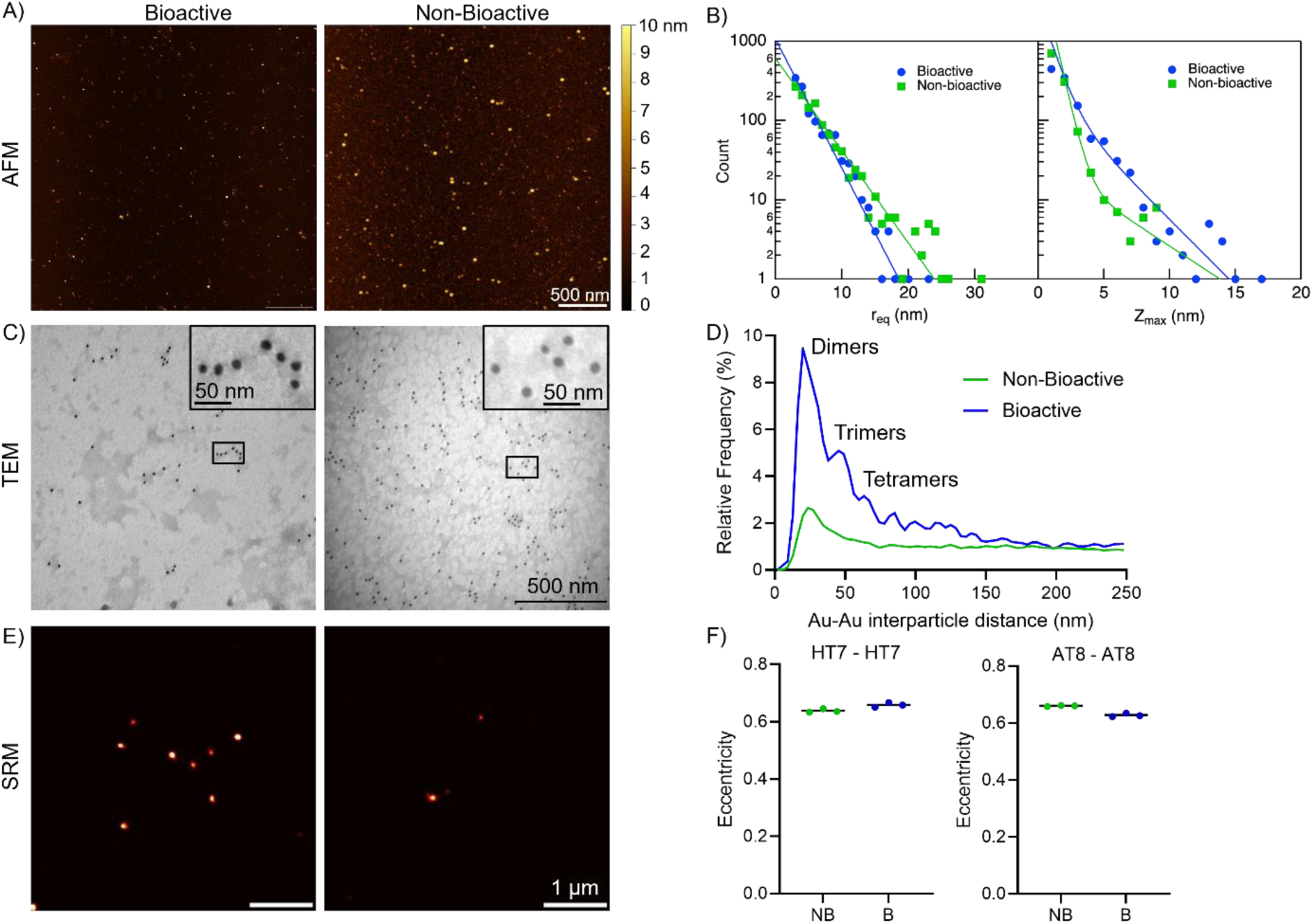
Non-bioactive and bioactive species contain small diffusible aggregates. Representative atomic force microscopy (AFM) images of bioactive and non-bioactive species isolated from the AD case 1971. B) AFM-based equivalent radius (r_eq_) and maximum height (Z_max_) distributions for non-bioactive (n = 1,141) and bioactive particles (n = 1,141), fit to single exponential and double exponential decaying functions, respectively. C) Representative transmission electron microscopy (TEM) immunogold images of bioactive and non-bioactive species labelled with the antibody pan-tau D5D8N and anti-rabbit secondary antibody conjugated with 12nm gold particles. Insets, representative gold particles pattern of non-bioactive and bioactive species. D) Analysis of the gold-gold distances from the center of particles. E) Representative super-resolution microscopy (SRM) images of bioactive and non-bioactive species captured and probed with AT8 antibody. F) Eccentricity quantification of HT7-HT7 and AT8-AT8 particles. Data represents mean of three individual measurements, conducted on non-bioactive and bioactive species isolated from the AD case 1971.

To further characterize these samples, we performed transmission electron microscopy (TEM) combined with tau-specific immunogold labeling. In both samples, immunogold confirmed the presence of tau clusters with no detectable fibrils (Figure 3C). Interestingly, the distance between gold particles reveals different patterns between the two species with higher number of dimers and linear trimers in the bioactive fraction, Au-Au distance of 22.5 and 45.0 nm, respectively, than in the non-bioactive tau samples (Figure 3D). Of note, we confirmed that the observed gold particles were due to tau presence rather than non-specific binding of secondary antibodies by omitting the primary anti-tau antibody, which resulted in a low number of gold particles. Moreover, analysis of the distances between gold particles revealed discrete values indicative of dimers, trimers, and tetramers in the bioactive species, which were not detected in the non-bioactive species (Figure 3). Tau observed across the micrographs did not display a uniform pattern, as would be expected for monomeric soluble tau derived from a recombinant source or human brain control lysate (Supplemental figure S6).

We also employed a recently developed single-molecule pull-down assay, which combines antibody-based immunoprecipitation with single-molecule super-resolution fluorescence microscopy, to analyze tau aggregates^18^. In this assay, tau aggregates are captured on an antibody-coated slide and detected using the same monoclonal antibody fluorescently labeled. In this assay, an eccentricity above 0.9 indicates the presence of fibrils^18^. Using this approach, we evaluated the eccentricity of both non-bioactive and bioactive species with super-resolution microscopy, probing with HT7 and AT8 antibodies. In both cases, eccentricity values were consistently below 0.9, indicating the presence of small amorphous aggregates.

Finally, upon fractionation of the non-bioactive and bioactive species by ultra-centrifugation 30 min at 100,000g at 4°C, we found that both species were present in the supernatant (Supplemental Figure S6). Notably, the bioactivity of the bioactive species was retained in the supernatant. This result is consistent with the presence of small diffusible aggregates and PBS soluble species. Overall, across three independent methods, we consistently found that both species are composed of morphologically similar, but biochemically distinct, small diffusible aggregates. While the negative stained EM images do not reveal fibrils, we cannot rule out the presence of rare, short fibrils, and refer to these structures therefore as small aggregates. Further studies including Cryo-EM would be necessary to obtain new insights and better characterized the nature of these aggregates.

### Non-bioactive and bioactive species differ in molecular conformations and phosphorylation status

We next further explored whether the bioactive and non-bioactive subfractions of the material eluting in the early fractions from size exclusion chromatography differ biochemically. We showed that these two species can be separated by anion exchange chromatography, suggesting differences in net charge or the exposure of charged regions. To explore these differences, we employed an ultrasensitive protein aggregate quantification assay based on single molecule array (SIMOA) technology^19^. This assay captures and detects tau aggregates using the same monoclonal antibody. Using the pan-tau antibody HT7, whose epitope is similar as D5D8N, we evaluated the Tau aggregate content and we found that both species contained aggregates, with higher levels in the bioactive species than in the non-bioactive species, 52.4 ng/mL and 27.4 ng/mL, respectively (Figure 4A). Interestingly, when we tested the MC1 antibody, developed to recognize a unique conformation of tau present in PHF as well as earlier stages^30^, it detected lower amounts of aggregated tau (∼5 ng/mL) than using the double monoclonal strategy in both species (Figure 4B). When we assessed phosphorylated aggregates using AT8, p181, and p217 antibodies, the bioactive species showed higher capture and detection signals. This indicates more exposed phosphorylation sites, which may explain the increased retention time on the AIEX column. Finally, we evaluated the phosphorylation profile of bioactive and non-bioactive Tau species from two AD cases using a combination of phospho-specific Tau antibodies in dot blot analyses (Figure 4B). For quantification, the phospho-Tau signal was normalized to the Tau signal detected by the pan-tau D5D8N antibody. Given that absolute signal intensities varied across experiments while the relative signal between the non-bioactive and bioactive species remained consistent, we compared, for each phosphosite, the phosphorylation signal ratio between the two species after normalization to the bioactive species. Raw signals and inter-experiment variability are shown in supplemental figures 7 and 8. Consistent with the SIMOA results, the dot blot quantification revealed that bioactive species are more phosphorylated than their non-bioactive counterparts (Figure 4C, D). Notably, for the AD case 1971 three antibodies targeting five phospho-sites pS202-pT205 (AT8), pT231, and pS396-pS404 (PHF1) showed higher immunoreactivity in the bioactive species. This increase in phosphorylation was even more pronounced for the bioactive species from AD case 2014, which displayed higher immunoreactivity than the non-bioactive species at all AD case 1971 phosphosites as well as for pT181 and pT231. These phosphorylation sites, located outside the AD fibril core region (amino acids 306–378), may contribute to the retention of these species on the anion exchange column. Using the same dot-blot quantification, we next estimated the relative phosphorylation burden of bioactive versus non-bioactive tau more globally. Across all phospho-epitopes examined, non-bioactive species ranged from 0.04 to 0.9 when expressed relative to the bioactive fraction (set to 1), corresponding to approximately 1- to 23-fold higher phospho-signals in the bioactive species (Supplementary Table 1). When phospho-epitopes were considered jointly, including the double-phospho antibodies AT8 and PHF1 but excluding the single-site pS396 and pS404 antibodies that interrogate the same region, the composite phospho-signal was increased by ∼4.5-fold in the bioactive fraction for AD case 1971 and by ∼7.8-fold for AD case 2014. Conversely, when we considered pS396 and pS404 but excluded PHF1, the composite fold-change was 4.3 for AD case 1971 and 9.4 for AD case 2014. The similar values for AD case 1971 are consistent with both pS396 and pS404 showing comparably lower signals in the non-bioactive fraction, whereas the larger fold-change for AD case 2014 in this analysis is mainly driven by pS404. In AD case 1971, pS396-only and pS404-only antibodies each show modest, non-significant individual changes, but tau molecules phosphorylated at both sites (PHF1 epitope), as well as at AT8 and pT217, are clearly enriched in the bioactive fraction. In AD case 2014, tau molecules carrying pS404 alone and/or double phosphorylation at S396/S404 are enriched in the bioactive fraction in addition to those recognized by pT181, AT8, pT217 and pT231. Together, these data indicate that, although both species are phosphorylated and one is globally more phosphorylated than the other, clusters of phospho-modifications—rather than a uniform increase at individual sites—are preferentially enriched in the bioactive fraction. This is consistent with the non-bioactive and bioactive tau species differing in their surface available charge, phosphorylation status, and the likelihood of containing (at least) homodimers of phosphorylated tau.

**Figure 4.**
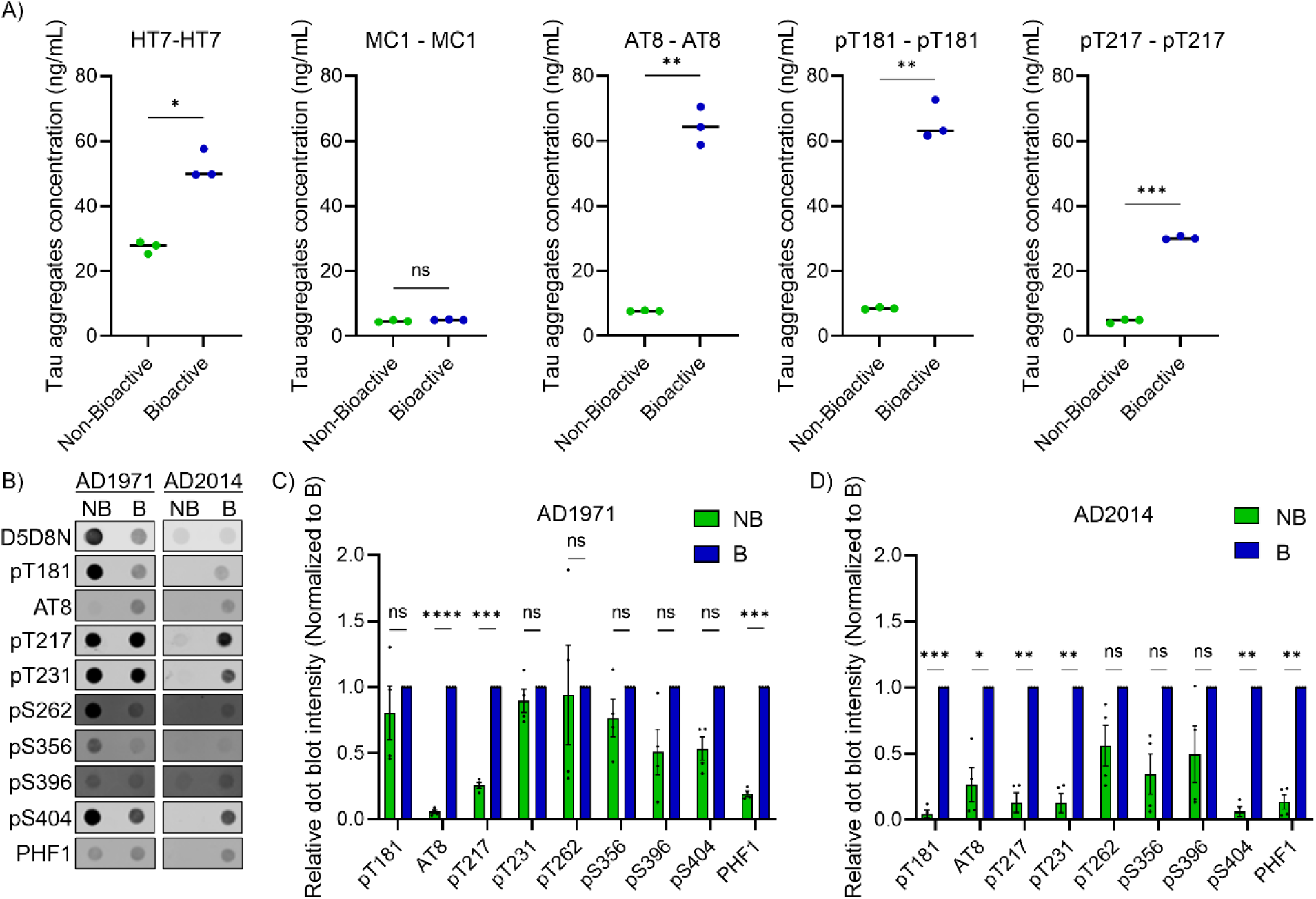
Tau Bioactive species exposes more phosphorylation sites. A) Tau aggregates concentration quantified by SIMOA assay using total tau antibody HT7, MC1, phospho-antibodies AT8, pT181 and pT217. Results were analyzed using the paired student t-test, * for p < 0.05, ** for p < 0.01, *** for p < 0.001, **** for p < 0.0001 and ns for non-significant. Experiments were conducted with non-bioactive and bioactive species extracted from AD case 1971. B) Dot blots of non-bioactive and bioactive species originating from two AD cases, probed with pan-Tau antibody D5D8Nalong with several anti-phospho Tau antibodies. C, D) Dot blots quantification, the anti-phospho Tau signal was adjusted to the signal obtain from the D5D8N pan-Tau antibody. Relative intensities of non-bioactive and bioactive species were determined from four independent experiments and normalized to the bioactive species (raw densitometry values are shown in figures S7 and S8). Non-bioactive and bioactive species are originating from two AD cases as indicated in panel B, C and D.

### Bioactive oligomeric tau is exceedingly potent

Based on the SIMOA experiment, we determined that the concentration of non-bioactive and bioactive species at 27.4 and 52.4 ng/mL, respectively, corresponding to approximately 12 ng of bioactive tau species per gram of frontal grey matter from a Braak VI case (i.e. pM concentrations). It is important to note, that these calculations were based on Tau species extracted from frontal cortex of Braak VI AD cases and hence Tau amounts might vary across brain regions and Braak stages. Given this low concentration, we investigated whether that low concentration could trigger aggregation in cells. Using the apparent molecular weight of HMW tau from SEC (∼500 kDa), we calculated concentrations of 55 pM and 105 pM for the non-bioactive and bioactive species, respectively. These samples were then used to treat FRET reporter cells at decreasing concentrations. It is important to note that the tau species were diluted 19-fold with media and lipofectamine, resulting in maximum concentrations of 2900 fM and 5600 fM for the non-bioactive and bioactive species, respectively. Starting from these initial concentrations, we performed a 7-point twofold dilution series, with seeding detected at concentrations as low as 175 fM. For this analysis, we focused solely on the FRET signal intensity, disregarding the number of FRET-positive cells, to determine the minimum tau seed concentration required to induce aggregation. The cell-based assay isolates misfolded templating from cell uptake or escape from endosomal and lysosomal pathways, providing an estimate of the amount of seed competent tau which is the lower limit for a positive signal.

The data of the number of seeded cells as a function of tau oligomer concentration, are fit well by a “one hit model”, i.e. a model in which a single seed is sufficient to induce daughter seeds in the reporter cells, consistent with predictions for prion-like behavior of the bioactive seeds (Fig 5B). We find that the seeding efficiency is more than 20-fold higher for the bioactive than the non-bioactive species, assuming the one hit model in both cases. This suggests that the bioactive seeds are at least an order of magnitude more potent than the non-bioactive seeds. It is important to note, however, that these calculations apply to a reporter system engineered to efficiently amplify seeds, with excess truncated tau reporter present in the cytoplasm, and access to the cytoplasm ensured by lipophilic reagents; in biological systems, the efficiency of seeding would be expected to be substantially less.

**Figure 5.**
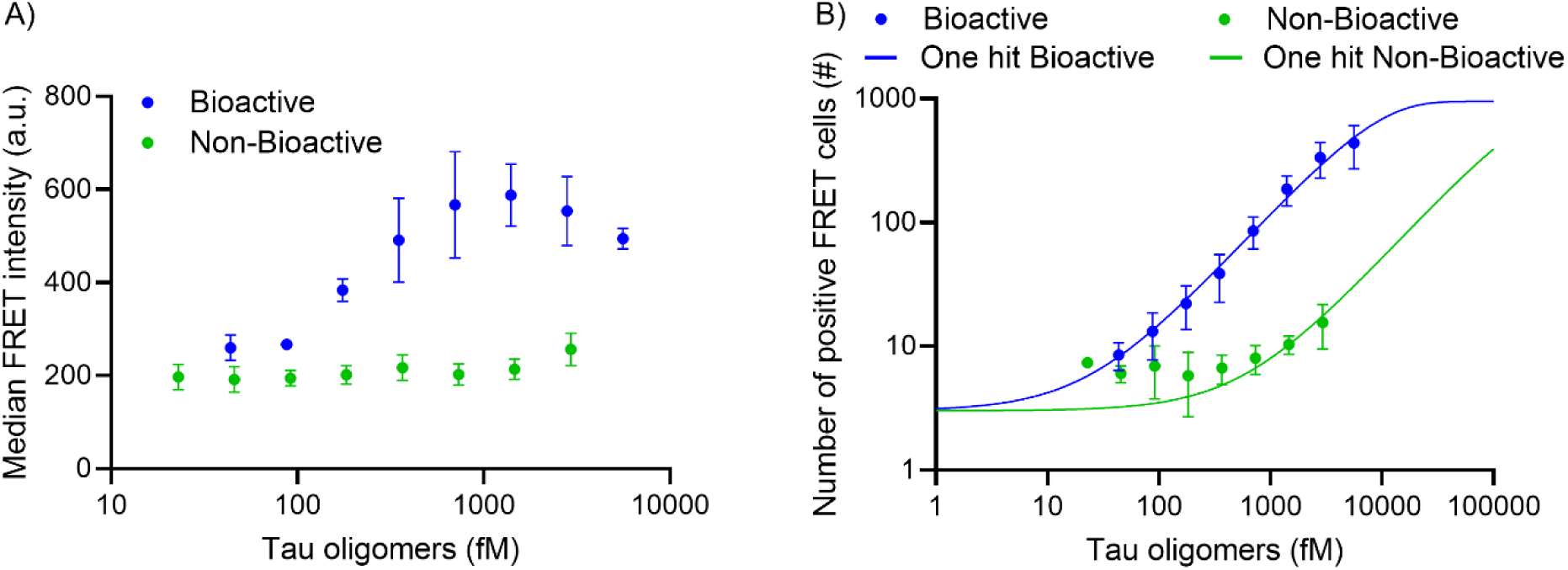
Seeding competency of bioactive tau. A) Data represents the median ± S.D. FRET intensity and B) number ± S.D. of seeded cells measured from three individual experiments performed in triplicate using the reporter HEK cell FRET assay with serially diluted samples. The initial concentrations were 2900 fM for the non-bioactive species (green) and 5600 fM for the bioactive species (blue), with each subsequent point representing a 1:2 dilution. Solid lines are fits to a “one hit” model as detailed in the Methods. Experiments were conducted with non-bioactive and bioactive species extracted from AD case 1971.

### Brain derived bioactive oligomeric tau species support RT-QuIC amplification of truncated tau and retain bioactivity in cellulo after amplification, unlike non-bioactive oligomers

In addition to assessing the high potency of the bioactive species in a cell reporter system, we also analyzed their ability to generate fibrils in an isolated biochemical assay. To this end, we performed Real-Time Quaking-Induced Conversion (RT-QuIC) assays to amplify and generate “daughter” bioactive seeds. We then tested whether these newly formed seeds retained their bioactivity through a second round of amplification via RT-QuIC. For the amplification, the AD core fragment (residues 306–378), which has been previously validated in RT-QuIC assays to produce a ThT-positive signal, reflecting fibril formation, when incubated with biological samples^20^, was used as a substrate (Figure 6A). Interestingly, only the bioactive species produced a detectable ThT signal consistent with the bioactive competency of these species observed *in cellulo* (Figure 6B). By TEM we confirmed the presence of fibrillar structures (Figure 6D). After 72 hours, ThT-free reaction mixtures were collected and subjected to a second amplification with fresh substrate. Similar to the first amplification, a ThT signal was observed only when daughter seeds were formed with bioactive species, although with a longer lag phase (11.4 hours vs. 15.8 hours; Figure 6B, C).

**Figure 6.**
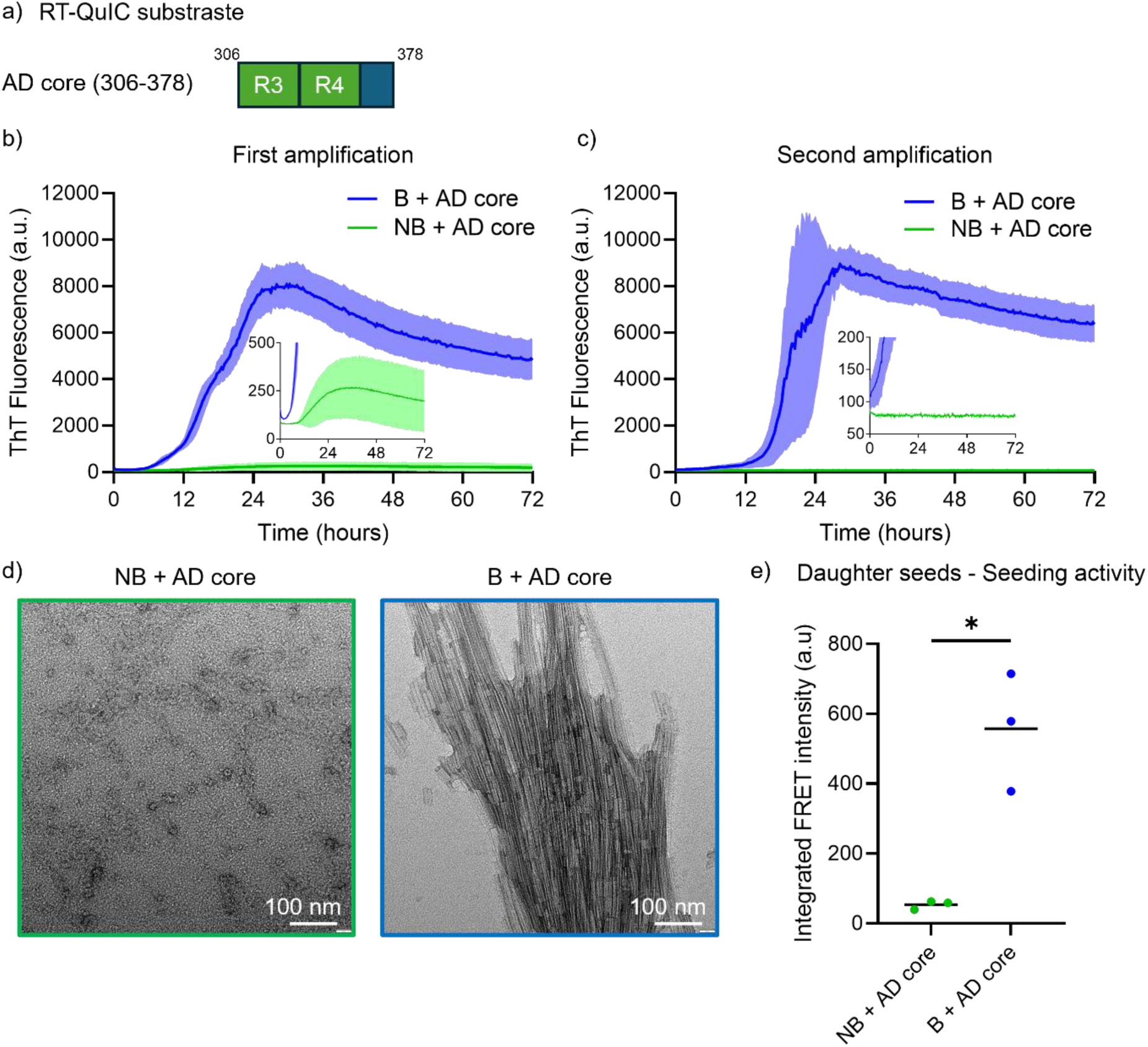
Bioactive species induce tau aggregation to generate seed competent daughters. A) Schematic representation of the AD core (residues 306-378) used as a substrate for RT-QuIC assays. A substrate concentration of 10 µM. B) RT-QuIC assays were performed by mixing AD core with non-bioactive (NB) and bioactive (B) species as seeds. C) RT-QuIC assays were performed with species formed after the 72 hours of incubation with both species and AD core acting as daughter seeds and fresh AD core. B, C) Data represents the mean ± S.D. of three individual experiments performed in triplicate. D) Representative negative-stain TEM images of daughter seeds formed with non-bioactive (NB) and bioactive (B) species after the first amplification (72h). E) Seeding activity of the daughter seeds (first amplification, 72h) is determined using the FRET reporter HEK cell assay and measured by flow cytometry. Data from three individual experiments are represented as mean, and analyzed using the paired student t-test, * for p < 0.05. Experiments were conducted on non-bioactive and bioactive species isolated from the AD case 1971.

Finally, we assessed the bioactivity of the daughter seeds generated after the first round of amplification *in cellulo*. Daughter seeds derived from the bioactive species retained bioactivity, whereas material generated from non-bioactive oligomers remained inactive, indicating that seeding-associated properties are preserved over amplification (Figure 6E). Interestingly, when bioactive species were incubated with a mixture of 0N3R and 0N4R tau isoforms, we did not observe an increased ThT signal, but we did detect small aggregates by TEM (Supplementary Figure S9). Moreover, the low seeding activity observed for the daughter seeds generated using the 0N isoform mixture may be attributed to the absence of phosphorylation sites on the recombinant substrates, as the 0N isoforms, expressed in *E. coli*, are not post-translationally modified. Consequently, despite being aggregated by the bioactive species, these unmodified substrates may lack the structural or biochemical features required to support efficient seeding. As expected, control lysate did not generate a ThT signal with any of the substrates; this was also true for AD core or biological samples alone, including those containing bioactive species (Supplemental Figure S10). Collectively, these results show that brain-derived bioactive tau oligomers can template the aggregation of a naïve tau fragment in vitro and propagate seeding activity across serial RT-QuIC amplification, with corresponding bioactivity in a cellular assay—consistent with a prion-like, templated misfolding mechanism.

## Discussion

This study aimed to isolate, enrich, and identify bioactive diffusible tau aggregates implicated in AD progression, providing insights into the biochemical features critical for tau seeding competency. Using 2D gels and *in vitro* dephosphorylation, we confirmed that HMW tau comprises multiple subspecies, with surface phosphorylated species identified as the bioactive subset, consistent with prior findings^16,22^.

Anion exchange chromatography of human derived material revealed the presence of a myriad of HMW tau species differing in charge and/or conformation. Given the extensive phosphorylation sites identified in soluble tau across multiple AD cases—more than 20^23,30,31^—it is unsurprising that various combinations might exist and yield multiple species. Nonetheless, this approach reproducibly separated non-bioactive and bioactive HMW tau species across nine AD cases. This finding aligns with studies showing that aggregates of the same protein can differ in biochemical properties and toxicity^32^, that monomeric tau may adopt inert and bioactive conformations in vitro^33^, and that tau aggregates derived from different tauopathies vary in conformation and bioactivity^34^. However, to our knowledge, this is the first demonstration that tau oligomers isolated from individual human AD brains can differ so drastically in biochemical properties and seeding competency.

The anion exchange chromatography data further suggests that the tau oligomers retained on the anion exchange column are the most likely to seed aggregates. This implies that specific intracellular environments or interactions may be critical for seeding. For instance, tau species with a pI above 7.4 would be positively charged in the neuronal cytoplasm, while those with a pI below 7.4 would carry an atypical negative charge for tau. We postulate that such heavily modified, negatively charged tau could recruit positively charged tau, triggering aggregation. This is reinforced by the data shown in Fig 4, illustrating that the bioactive fraction has phosphorylation epitopes detectable by dot blots and SIMOA assays that detect multimers that are multiply phosphorylated. Indeed, using available anti-Tau phosphoepitopes we showed that bioactive species were phosphorylated at positions T181, S202, T205, T217, T231, S262, S356, S396 and S404 with stronger immunoreactivity for phosphosites T217, T231 and S404 comparatively to the non-bioactive species, which could suggest that these epitopes are available for interaction with the anion exchange column. While we cannot yet assign specific phospho-epitopes or define a phosphorylation signature for seeding activity, our data indicate that phosphorylation clusters are enriched in the bioactive fraction and seeding competence requires tau that is both assembled and highly phosphorylated. TEM and AFM are label-free high-resolution methods, which rely on the drying of the sample on grids or mica. The affinity to small and large aggregates varies, which causes higher or lower average values of the diameter of sampled species compared to the distribution of the populations in solution. Conversely, single-molecule fluorescence is insensitive to single monomers and capable of measuring the size of very large aggregates at a scale of 200 × 200 µm skewing the average to larger values. Thus, although the exact measurements are somewhat technique dependent, all three techniques used converge on the finding that the non-bioactive and bioactive species are similar but not identical small aggregates that are nonfibrillar.

Combining these observations refines the characteristics that define seeding-competent tau. In our preparations, both fractions consist of small, non-fibrillar oligomers, yet only the highly phosphorylated bioactive oligomers are strongly seed-competent, whereas non-bioactive oligomers of comparable size show little or no seeding activity. This supports the view that size or fibrillar status per se is not the primary determinant of bioactivity and that phosphorylation acts as a key gating factor. This interpretation is consistent with work on recombinant hyperphosphorylated tau, where multi-site phosphorylation is sufficient to drive spontaneous aggregation into small, predominantly non-fibrillar oligomers that are more toxic and more efficient at nucleating unmodified tau than unphosphorylated tau ^35^. Recent single-molecule AFM studies further show that hyperphosphorylation stabilizes smaller, stiffer and more adhesive tau oligomers, supporting a model in which phosphorylation reshapes both the structural and mechanical landscape of early aggregates in ways that favour pathogenic seeding^36^. Finally, studies using AD-derived tau filaments have demonstrated that the fibril core structure alone is insufficient to recapitulate the seeding potency of brain-derived material and that phosphorylation within the disordered fuzzy coat is required for full seeding competency in neurons and *in vivo*^37^. Together, these findings and our own data converge on the idea that tau seeding competence is best understood as a property of a restricted subset of aggregated species that are both assembled and decorated with clustered phospho-epitopes, rather than a generic feature of all oligomers or fibrils.

Our findings also demonstrate that tau recruitment and aggregation can occur at extremely low concentrations of tau oligomers in the cytoplasm, where they encounter naïve tau. However, this bypasses processes such as tau uptake and endolysosomal escape, which might necessitate higher concentrations, as previously suggested^38^. Moreover, we showed that the small diffusible aggregated bioactive species induce the formation of bioactive daughter seeds conversely to sarkosyl extracted fibrillar tau^39^. The low concentration of seeds may explain the difficulties in developing a RT-QuIC assay for tau seeds in biofluids.

In previous studies we demonstrated by mass spectrometry that both the HMW fraction, (as well as an MC1 (conformationally specific tau antibody) immunoprecipitation fraction), contain a subset of the post translational modifications that are present on sarkosyl insoluble fibrillar tau in AD^23,30^, suggesting that the HMW fraction is a form fruste of the fibrillar fraction. On the other hand, injection of HMW tau into h-tau transgenic mice that express full length 3R and 4R tau^40^, showed that the HMW fraction propagated across neural systems but did not mature into sarkosyl insoluble fibrils over the course of 3 months. Our current results suggest a possible way to resolve this discrepancy: in this study, truncated tau, rather than a mix of 3R/4R recombinant full length tau, supports fibril formation in the RT-QuIC assay, suggesting that the conformation of the templated misfolding product is at least in part consequent to the specific substrate; these results are broadly compatible with the cryo-EM studies of Lovestäm *et al.*, 2022^41^, which demonstrated that extending the amino acid sequence of recombinant tau beyond the core 297-391 fragment impaired or dramatically altered fibril formation. These results also bring to mind the surprisingly sudden appearance of tau fibrils in vivo (as observed by in vivo multiphoton microscopy) after protease activation and appearance of cleaved tau^42^, perhaps explaining the surprising and puzzling observation that the extremely stable β pleated sheet containing tau fibrils can appear so quickly.

In the prion field, a “seed” is defined as a misfolded conformer of the prion protein capable of inducing the misfolding of native prion molecules, with daughter seeds retaining the ability to propagate aggregation and spread throughout the brain^43^. Owing to the parallels between prion propagation and tau aggregation, it has been hypothesized that tau may spread via a prion-like mechanism^3^. Supporting this notion, it has been demonstrated that tau seeds formed *in cellulo* can replicate and retain seeding competency over multiple passages^44,45^. However, contrasting results have shown that Sarkosyl-insoluble extracts from Alzheimer’s disease (AD) brains can induce tau aggregation *in vitro*, yet the resulting daughter seeds may lack sustained seeding competency^39^. These findings suggest that the mechanisms underlying tau seeding remain incompletely understood—particularly regarding the precise identity of the tau species that initiate and sustain this process. In this context, our fractionation and orthogonal imaging analyses indicate that seed competence is not explained by an oligomeric form alone: both bioactive and non-bioactive HMW tau are predominantly small assemblies (dimers–tetramers), yet only the bioactive subset retains robust activity in vitro and in cellulo. This points to a two-component requirement for bioactivity, whereby tau must be assembled into an aggregated state and must also carry specific biochemical attributes—most notably features enriched in the anion-exchange–retained fraction and correlated with surface-accessible phosphorylation—that distinguish seed-competent from seed-incompetent oligomers. Moreover, our current data suggest that HMW bioactive seeds can induce seeding in reporter cells linearly related to concentration (Fig 5B), consistent with a prion like behavior. It is important to note that the reporter cells have been designed to maximize seeding efficiency, using lipofectamine to bypass uptake mechanisms and an optimized truncated construct of the repeat domains to maximize aggregation. Thus, the current data supports the biochemical idea of templated misfolding at very low concentrations, but do not speak to the dose relevant for biological activity in an organism.

From a therapeutic perspective, effective tau-targeted therapies remain lacking^46^. In this study, we established a protocol to extract small diffusible aggregates, stable forms of tau capable of inducing aggregation, providing a foundation for developing reagents specifically targeting bioactive tau seeds in AD.

## Acknowledgement

We acknowledge the use of the AFM instrument from the laboratory of Mark C. Williams at Northeastern University and Dr. Mike Morse from Williams Lab for assistance with the AFM.

## Authors contributions

NQ, BTH, and DK conceived the study and designed the experiments. NQ, DS, DB, YC, JEC, AW, VD, MBMSM, and FAB performed experiments and analyzed data. GM, MC and DGA developed mathematical models for seeding and TEM analyses. TC, AM, MF and DO selected and provided human brain tissues. The manuscript was written by NQ and BTH with input from all the authors.

