## Supplemental figures for "Rare bioactive diffusible tau species from Alzheimer brain support both templated misfolding and fibril formation"

**Authors Affiliations:**

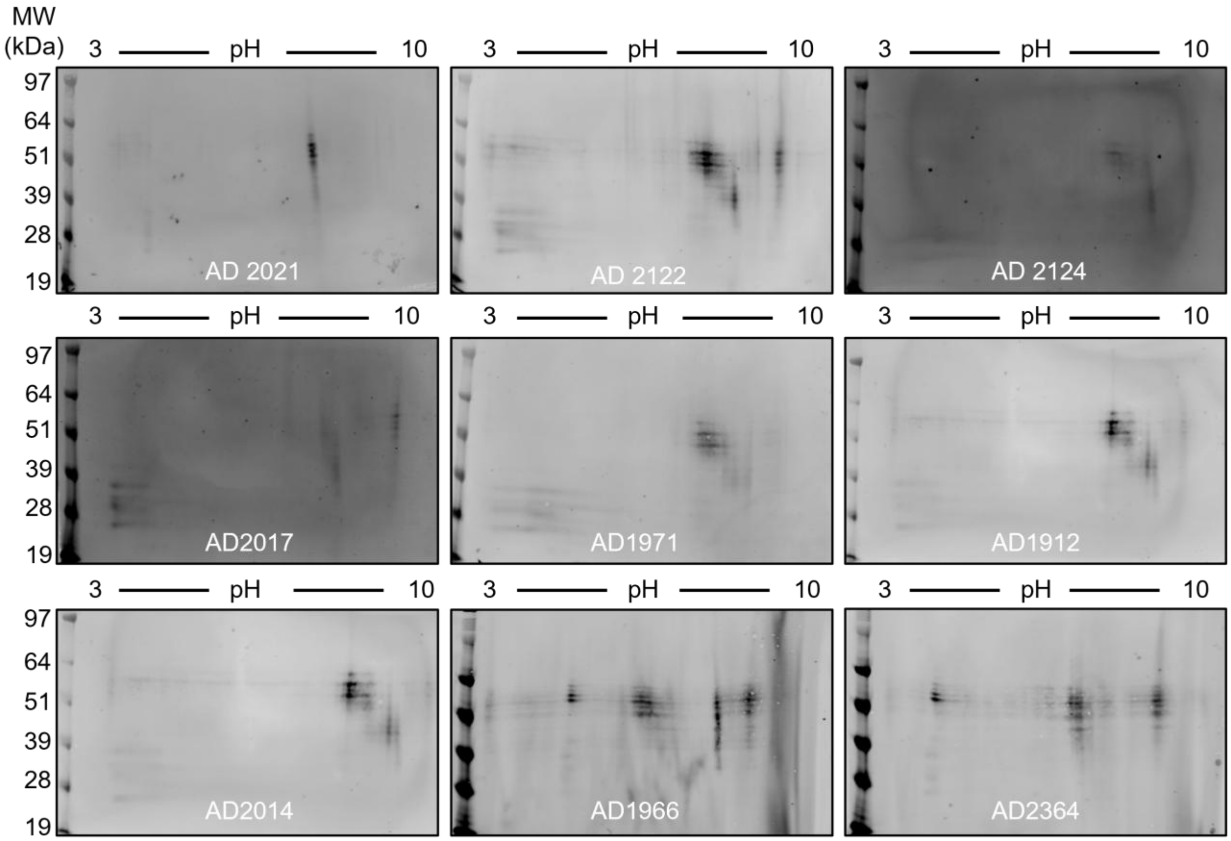


***Supplementary Figure S1. Tau molecules heterogeneity.*** *Two-dimensional gel electrophoresis of HMW Tau extracted from 9 AD cases, probed with D5D8N.*

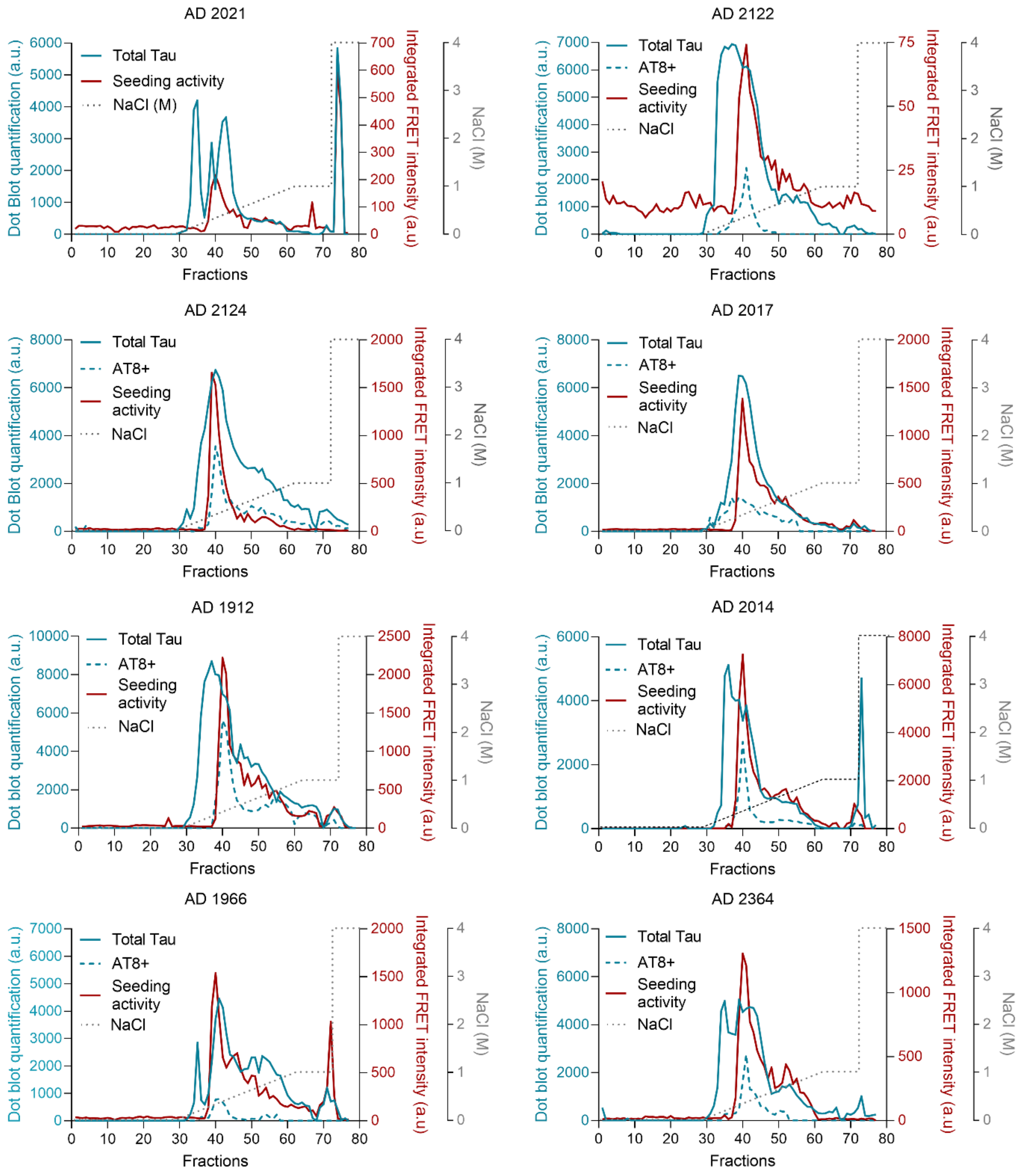


***Supplementary Figure S2. HMW Tau non-bioactive and bioactive species found across AD cases.*** *Fractionation of HMW Tau population derived from 9 human AD brains via anion exchange chromatography (AIEX) and elution gradient of increasing NaCl concentration (dashed line). Tau presence in each fraction is evaluated by dot blot probed with D5D8N (blue line) and phosphorylated Tau is evaluated with AT8 antibody (red line). Seeding activity of each fraction is determined with the FRET reporter HEK cell assay and measured by flow cytometry (green line), data from one seeding assay performed in triplicate, out of three, is plotted as representative of the bioactivity profile.*

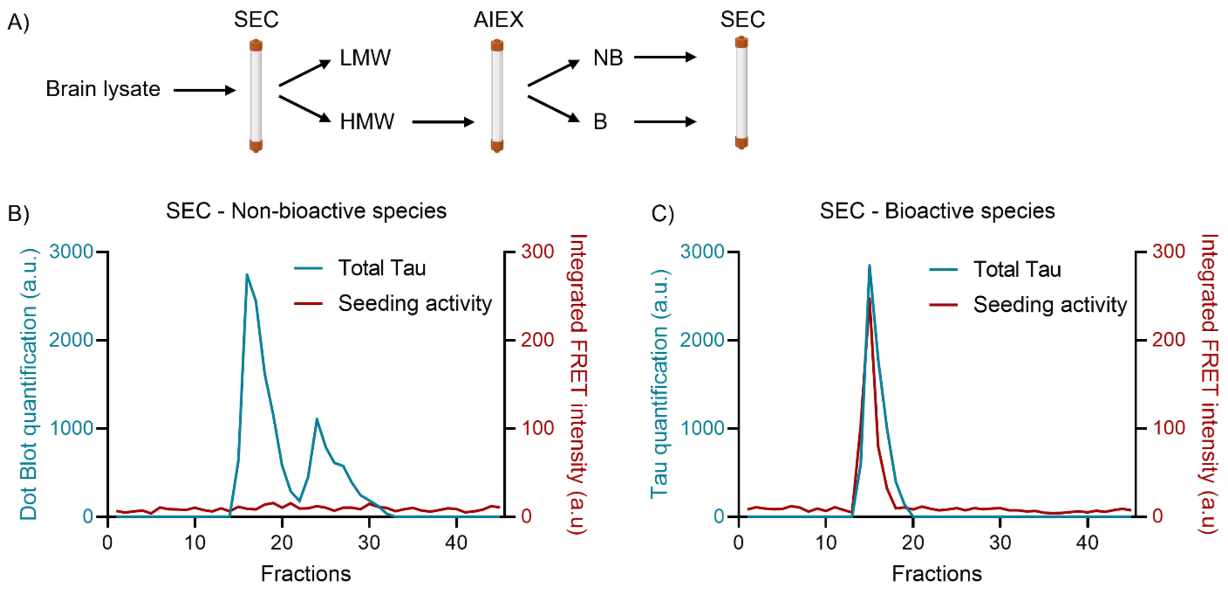


***Supplementary Figure S3.*** *A) Validation via size exclusion chromatography of non-bioactive and bioactive species maintaining their size following sequential chromatographic strategies. SEC: Size Exclusion Chromatography, AIEX: Anion Exchange Chromatography, LMW: Low Molecular Weight Tau, HMW: High Molecular Weight Tau, NB: Non-bioactive species, B: Bioactive species. B), C) Injection of non-bioactive and bioactive species onto a size exclusion column. Total Tau is measured in each fraction by dot blot quantification using D5D8N antibody (blue line). Seeding activity of each fraction is determined using the FRET reporter HEK cell assay and measured by flow cytometry (green line), data from one seeding assay performed in triplicate, out of three, is plotted as representative of the bioactivity profile. Non-bioactive and bioactive species are isolated from AD case 1971.*

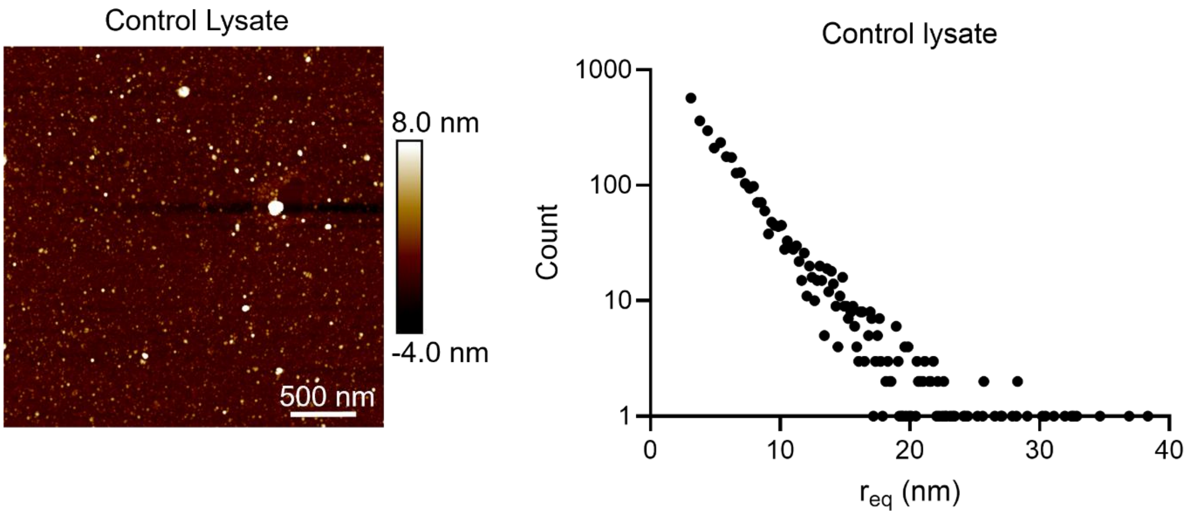


***Supplementary Figure S4.*** *Representative atomic force microscopy (AFM) images and equivalent radius ($r_{eq}$) distributions of control lysate (n=3,642).*

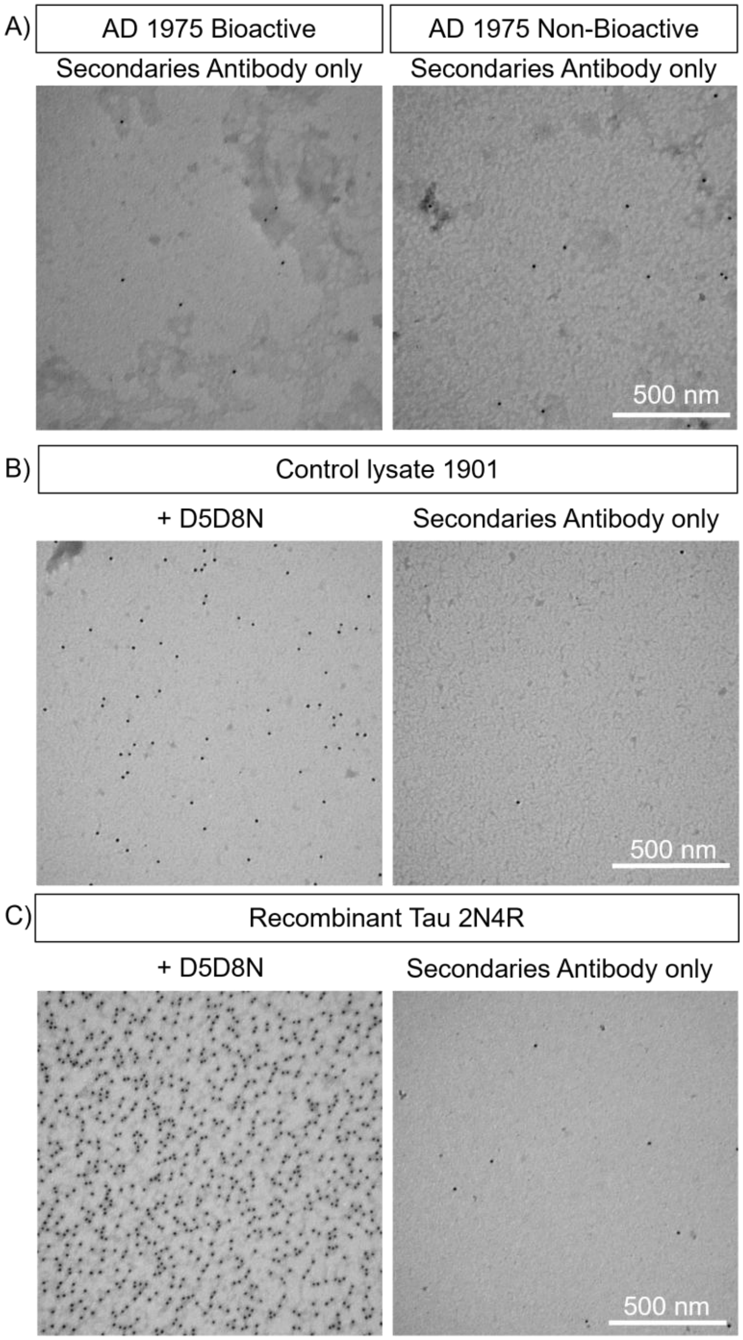


***Supplementary Figure S5.*** *Representative transmission electron microscopy (TEM) immunogold images of A) bioactive and non-bioactive species omitting primary antibody and incubated with an anti-rabbit secondary antibody conjugated with 12nm gold particles; B) Human brain control lysate in the presence or absence of the total Tau antibody D5D8N; C) Recombinant Tau 2N4R in presence or absence of the total Tau antibody D5D8N.*

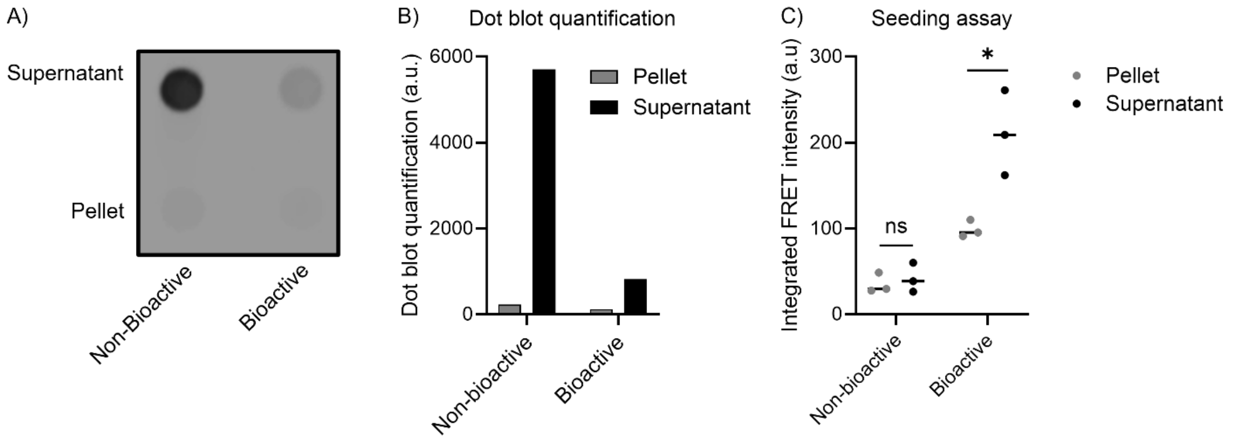


***Supplementary Figure S6.*** *Non-bioactive and bioactive species are fractionated via ultra-centrifugation. A) Dot blot of pellet and supernatant fractions of non-bioactive and bioactive species probed with the total Tau antibody D5D8N. B) Dot blot quantification. C) Seeding activity assay measured by flow cytometry. Data represents the mean ± S.D. of three individual experiments performed in triplicate. Non-bioactive and bioactive species are isolated from AD case 1971.*

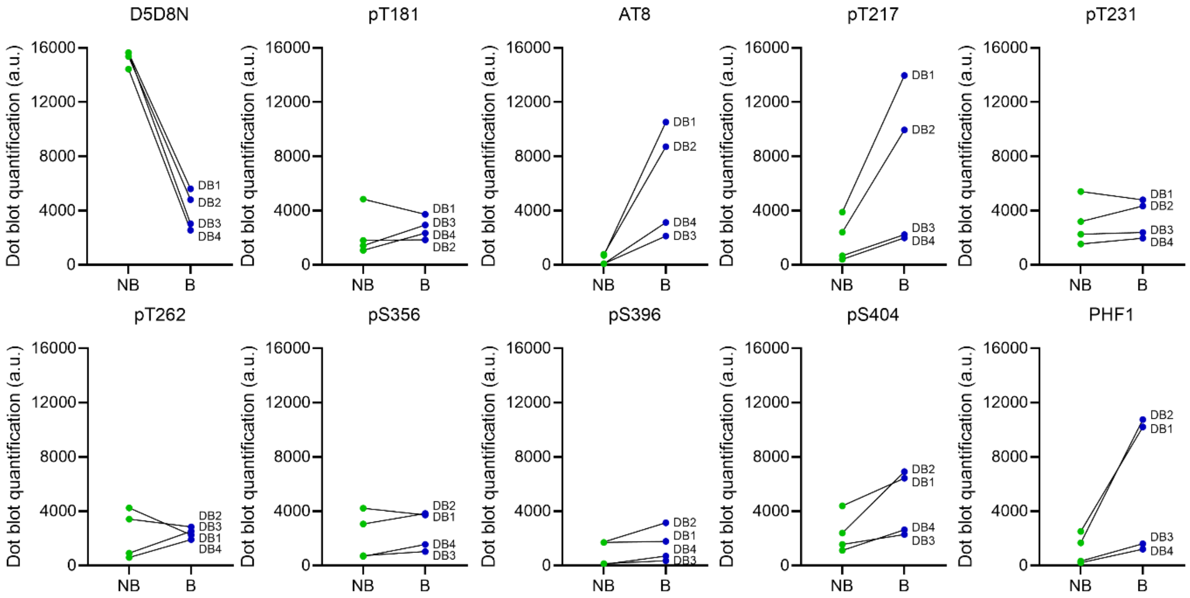


***Supplementary Figure S7.*** *Dot blot quantification of four independent experiments (DB1, DB2, DB3 and DB4) probed with pan Tau antibody D5D8N and anti-phospho Tau antibodies as indicated on corresponding panels. Non-bioactive and bioactive species are originating from the AD case 1971.*

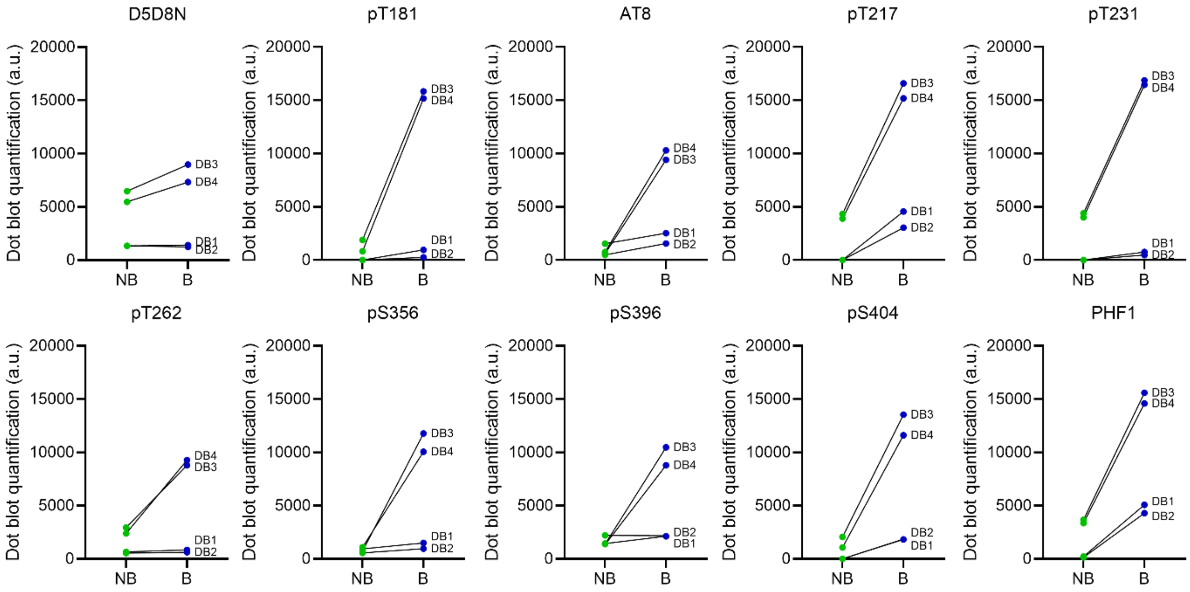


***Supplementary Figure S8.*** *Dot blot quantification of four independent experiments (DB1, DB2, DB3 and DB4) probed with pan Tau antibody D5D8N and anti-phospho Tau antibodies as indicated on corresponding panels. Non-bioactive and bioactive species are originating from the AD case 2014.*

***Supplementary table 1.*** *Fold change of phospho-epitope signal in bioactive versus non-bioactive species. A value of 1 indicates no difference, whereas values greater than 1 indicate higher phosphorylation in the bioactive species. Asterisks denote phospho-epitopes for which the difference between non-bioactive and bioactive species was statistically significant for each case, based on the dot blot quantification shown in Figure 4.*

| | pT181 | AT8* | pT217* | pT231 | pT262 | pS356 | pS396 | pS404 | PHF1* |
|---|---|---|---|---|---|---|---|---|---|
| AD1971 | 1.2 | 17.6 | 3.9 | 1.1 | 1.1 | 1.3 | 2.0 | 1.9 | 5.2 |

| | pT181* | AT8* | pT217* | pT231* | pT262 | pS356 | pS396 | pS404* | PHF1* |
|---|---|---|---|---|---|---|---|---|---|
| AD2014 | 23.0 | 3.8 | 7.7 | 7.9 | 1.8 | 2.9 | 2.0 | 16.4 | 7.4 |

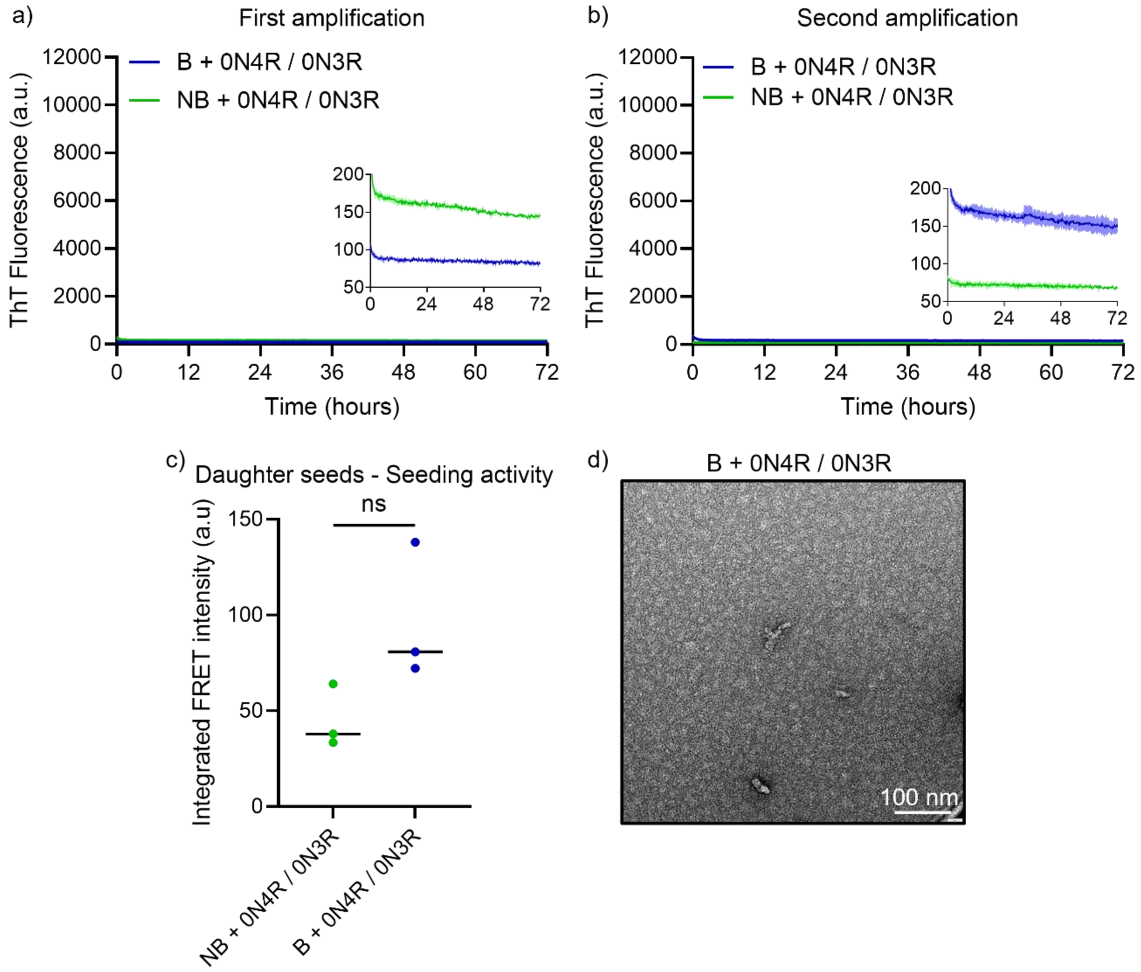


***Supplementary Figure S9.*** *RT-QuIC assays performed with Bioactive and Non-bioactive species isolated from AD case 1971. 0N4R / 0N3R isoforms were mix at a 2:3 ratio. A protein concentration of 10 µM was used for the total mixture of substrates. A) First amplification. B) Second amplification with samples from first amplification and substrates freshly prepared. Data represents the mean ± S.D. of three individual experiments performed in triplicate. C) Seeding activity of the daughter seeds (first amplification, 72h) is determined using the FRET reporter HEK cell assay and measured by flow cytometry. Data from three individual experiments are represented as mean, and analyzed using the paired student t-test, * for p < 0.05. D) Representative negative-stain TEM images of daughter seeds formed with bioactive (B) species and 0N isoforms after the first amplification (72h).*

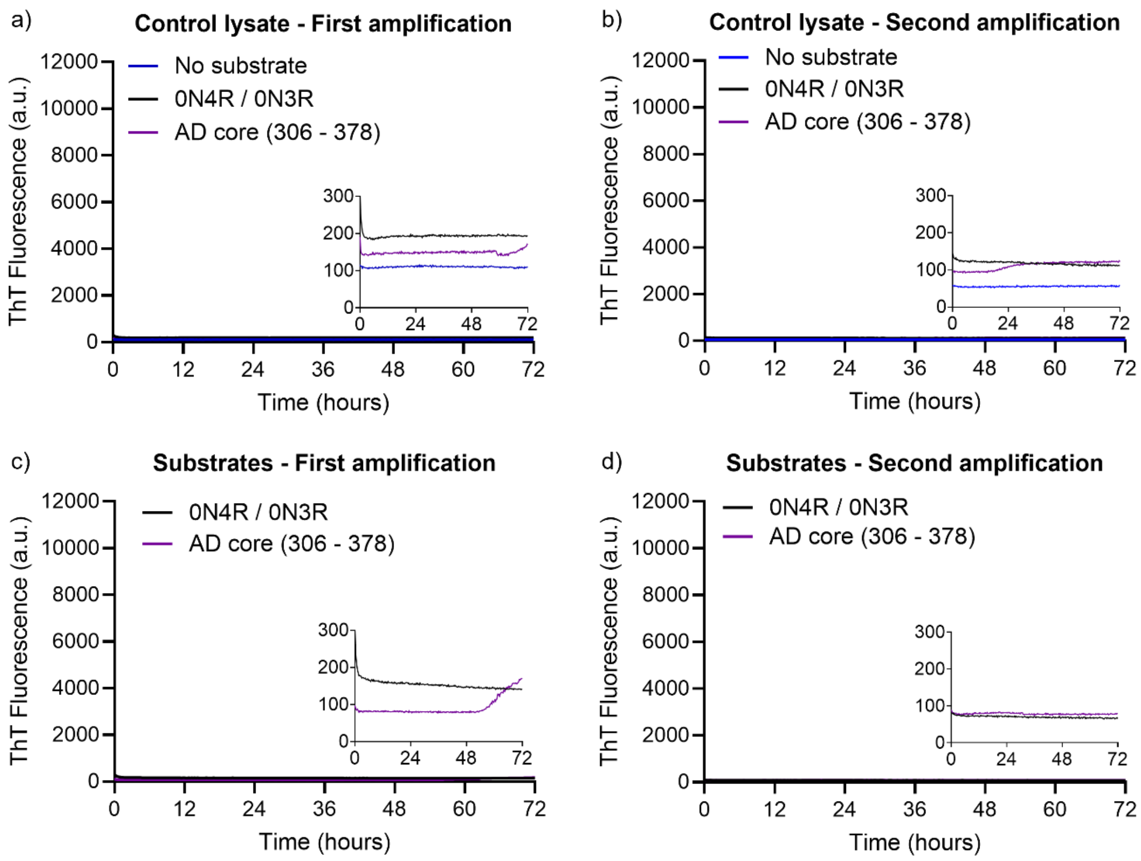


***Supplementary Figure S10.*** *RT-QuIC assays performed with control lysate and substrates without biological samples. 0N4R / 0N3R isoforms were mix at a 2:3 ratio and AD core (residues 306-378) was used a single substrate. A protein concentration of 10 µM was used for the two substrates. A), C), First amplification. B), D),) Second amplification with samples from first amplification and substrates freshly prepared. Data represents the mean ± S.D. of three individual experiments performed in triplicate.*
